# A 3.5-minute-long reading-based fMRI localizer for the language network

**DOI:** 10.1101/2024.07.02.601683

**Authors:** Greta Tuckute, Elizabeth Jiachen Lee, Aalok Sathe, Evelina Fedorenko

**Affiliations:** Department of Brain and Cognitive Sciences and McGovern Institute for Brain Research, Massachusetts Institute of Technology, Cambridge, Massachusetts, USA; Kempner Institute for the Study of Natural and Artificial Intelligence at Harvard University, Boston, Massachusetts, USA

## Abstract

The field of human cognitive neuroscience is increasingly acknowledging inter-individual differences in the precise locations of functional areas and the corresponding need for individual-level analyses in fMRI studies. One approach to identifying functional areas and networks within individual brains is based on robust and extensively validated ‘localizer’ paradigms—contrasts of conditions that aim to isolate some mental process of interest. Here, we present a new version of a localizer for the fronto-temporal language-selective network. This localizer is similar to a commonly-used localizer based on the reading of sentences and nonword sequences (Fedorenko et al., 2010) but uses speeded presentation (200ms per word/nonword). Based on a direct comparison between the standard version (450ms per word/nonword) and the speeded versions of the language localizer in 24 participants, we show that a single run of the speeded localizer (3.5 min) is highly effective at identifying the language-selective areas: indeed, it is more effective than the standard localizer given that it leads to an increased response to the critical (sentence) condition and a decreased response to the control (nonwords) condition. This localizer may therefore become the version of choice for identifying the language network in neurotypical adults or special populations (as long as they are proficient readers), especially when time is of essence.

## Introduction

Neuroscientific studies of uniquely human abilities rely predominantly on non-invasive neuroimaging techniques such as functional magnetic resonance imaging (fMRI). A widespread methodological approach in fMRI studies of human brain function is to average individual activation maps for some contrast of interest in a template brain space and perform statistical analyses in each voxel across individuals to derive a group-level whole-brain statistical map. However, functional regions vary in their precise locations across individuals (Fischl et al., 2008; Frost & Goebel, 2012; Tahmasebi et al., 2012; Vázquez-Rodríguez et al., 2019; Somers et al., 2021). Correspondingly, reliance on these group-averaging approaches can lead to low sensitivity and functional resolution (Brett et al., 2002; Saxe et al., 2006; Nieto-Castañón & Fedorenko, 2012). Inter-individual variability is particularly problematic when functional regions of interest lie in proximity to functionally distinct regions, as is the case with both frontal and temporal language regions (e.g., Tomaiuolo et al., 1999; Fedorenko et al., 2012; Tahmasebi et al., 2012; Deen et al., 2015; Braga et al., 2020; Du et al., 2024; see Fedorenko & Blank, 2020, for discussion of this issue for ‘Broca’s area’).

One increasingly popular solution that circumvents inter-individual variability in the precise locations of functional regions is the use of functional ‘localizers’ (Saxe et al., 2006; Nieto-Castañón & Fedorenko, 2012; Gratton & Braga, 2021). In this approach, a brain region or network that supports a mental process of interest is identified with a functional contrast in each individual brain and subsequently, the region’s/network’s responses to some critical condition(s) of interest are examined. Consistent use of these localizers across studies and labs (and in some cases, species; Russ et al., 2021) affords greater confidence that the ‘same’ region or set of regions is being studied, compared to relying on anatomical landmarks alone, and thus facilitates the accumulation of scientific knowledge.

The functional localization approach has been successful across many domains of perception and cognition including high-level visual and auditory processing, social cognition, and language (Kanwisher et al., 1997; Epstein & Kanwisher, 1998; Downing et al., 2001; Belin et al., 2002; Saxe & Kanwisher, 2003; Baker et al., 2007; Fedorenko et al., 2010, 2013; Overath et al., 2015; Fischer et al., 2016; Isik et al., 2017). In the domain of language, Fedorenko et al. (2010) developed a localizer that relies on a contrast between language processing and the processing of a perceptually similar condition that lacks linguistic structure or meaning (e.g., reading or listening to sentences vs. nonword lists, or listening to sentences vs. backwards speech or acoustically degraded sentences; Bedny et al., 2011; Scott et al., 2017; Lipkin et al., 2022; Malik-Moraleda, Ayyash et al., 2022). Such contrasts target brain areas that support computations related to accessing words and combining them into complex linguistic structures and meanings. These ‘language localizers’ robustly identify the left-lateralized fronto-temporal language network, which has long been implicated in language processing based on investigations of patients with aphasia (e.g., Luria, 1970; Goodglass, 1993; Bates et al., 2003; Fridriksson et al., 2018; Wilson et al., 2023) and group-averaging neuroimaging investigations of language processing (e.g., Binder et al., 1997; Price, 2010; Friederici, 2012). Importantly, language localizers are highly generalizable, eliciting similar activations across presentation modalities, materials, and tasks (see Fedorenko et al., 2024). Moreover, the brain regions that this localizer identifies closely correspond to those that emerge from the bottom-up clustering of voxel time-courses obtained during rest (Braga et al., 2020) or while performing tasks (Du et al., 2025; Shain & Fedorenko, 2025). This correspondence highlights that the language network is a ‘natural kind’ in the brain: an ontologically meaningful grouping of a set of brain regions that show highly synchronized activity over time. Neuroimaging studies of language that rely on the functional localization approach have produced a number of robust and replicable findings both about i) the relationship between language and other perceptual and cognitive processes (e.g., Fedorenko et al., 2011; Deen et al., 2015; Amalric & Dehaene, 2019; Ivanova et al., 2020; Jouravlev et al., 2019; Chen et al., 2023; Shain et al., 2023), and ii) the internal organization of the language system and its computations (e.g., Blank et al., 2016; Fedorenko et al., 2020; Shain et al., 2022; Shain, Kean et al., 2024).

One practical concern that researchers often express about the use of localizers is that they take time. Time is often a precious commodity in neuroimaging research, either because the critical task is already long and/or because the population of interest may have low tolerance for the scanner environment. However, given the advantages that localizers provide—including greater sensitivity, greater functional resolution, more accurate effect size estimation, higher interpretability of the responses in the critical tasks, and the ability to meaningfully accumulate knowledge across studies, labs, and species—many researchers continue to adopt this approach. One recent effort in the field has therefore been to try to optimize localizers so that they can be as short as possible while still yielding robust individual-level responses (e.g., Dodell-Feder et al., 2011; Lee et al., 2024; Marvi, Hutchinson et al., 2025).

In this study, we develop a shorter version of a widely used reading-based language localizer (Fedorenko et al., 2010). We leverage the fact that humans can read at fast rates, especially when the need for eye movements is minimized by presenting words one at a time in the center of the screen in a rapid serial visual presentation (RSVP) paradigm (e.g., Forster, 1970; Potter et al., 1980, 1986; Potter, 2012; Mollica & Piantadosi, 2017). In these studies, participants can process linguistic information even when each word is presented for as little as ∼80-200ms, as evidenced by accurate recall of the stimuli and high accuracy in answering comprehension questions about the content. Moreover, a few studies (Vagharchakian et al., 2012; Benjamin & Gaab, 2012; Christodoulou et al., 2014) have found that speeded reading, similar to reading at slower speeds, activates the language areas, but these studies have used a group-averaging approach, leaving open the question of whether speeded reading elicits sufficiently robust responses in individual participants. This is the question our study aims to address. Although this question is primarily methodological in nature, our study’s design allows us to additionally ask a theoretically interesting question about whether the increased processing difficulty due to speeded presentation affects neural responses in the language-selective network, or instead (or in addition) in the domain-general Multiple Demand network, which is sensitive to cognitive effort across diverse paradigms (e.g., Fedorenko et al., 2013; Duncan et al., 2012; Duncan, 2010; Duncan et al., 2020; Assem et al., 2020b).

## Methods

### Brief overview

24 adults each completed two versions of a language localizer task. In both versions, participants read sentences and lists of unconnected pronounceable nonwords presented on the screen one word/nonword at a time. The two versions differed in the presentation speed of each word/nonword. One localizer version was an extensively validated language localizer task (Fedorenko et al., 2010; Mahowald & Fedorenko, 2016; see Lipkin et al., 2022 for data from >600 participants on this version) where each word/nonword is presented for 450 ms (‘standard language localizer’). The other version was a new, speeded version of the task where each word/nonword was presented for 200 ms (‘speeded language localizer’). 22 of the 24 participants completed the two versions of the language localizer in the same scanning session; the remaining two—in separate sessions (1 and 463 days apart). For all participants, the speeded version was run after the standard version. Each scanning session lasted between 1 and 2 hours and included a variety of additional tasks for unrelated studies. The materials, scripts, and screen recordings for the two language localizer versions are available at https://www.evlab.mit.edu/resources-all/download-localizer-tasks (standard version) and https://github.com/el849/speeded_language_localizer/ (speeded version).

### Participants

24 neurotypical adults (12 female, 12 male), aged 18 to 60 (mean: 28.04; std: 9.25), participated for payment between June 2021 and December 2022. All participants were native speakers of English, had normal or corrected-to-normal vision, and no history of neurological, developmental, or language impairments. 22 participants (∼92%) were right-handed, as determined by the Edinburgh handedness inventory (Oldfield, 1971), 2 participants (∼8%) were left-handed. All participants gave informed written consent in accordance with the requirements of the MIT’s Committee on the Use of Humans as Experimental Subjects (COUHES).

### fMRI tasks

#### Language network localizer tasks

##### Standard language localizer task

A reading task contrasted *sentences* (e.g., THE SPEECH THAT THE POLITICIAN PREPARED WAS TOO LONG FOR THE MEETING) and lists of unconnected, pronounceable *nonwords* (e.g., LAS TUPING CUSARISTS FICK PRELL PRONT CRE POME VILLPA OLP WORNETIST CHO) in a standard blocked design with a counterbalanced condition order across runs, as introduced in Fedorenko et al. (2010). Each stimulus consisted of 12 words/nonwords. Stimuli were presented in the center of the screen, one word/nonword at a time, at the rate of 450 ms per word/nonword. Each stimulus was preceded by a 100 ms blank screen and followed by a 400 ms screen showing a picture of a finger pressing a button, and a blank screen for another 100 ms, for a total trial duration of 6 s. Participants were instructed to read attentively (silently, to themselves) and to press a button on the button box whenever they saw the picture of a finger pressing a button on the screen. The button-pressing task was included to help participants remain alert. Experimental blocks lasted 18 s (with 3 trials per block) and fixation blocks lasted 14 s. Each run (consisting of 16 experimental blocks and 5 fixation blocks) lasted 358 s (5 min 58 s). Participants completed 2 runs.

##### Speeded language localizer task

The speeded version of the language localizer was identical to the standard version except that each word/nonword was presented for 200 ms instead of 450 ms (i.e., ∼56% faster). Each stimulus was preceded by a 100 ms blank screen and followed by a 400 ms screen showing a picture of a finger pressing a button, and a blank screen for another 100 ms, for a total trial duration of 3 s. The instructions to the participants were the same as in the standard version although they were warned that the presentation would be somewhat fast, and they were told not to worry if they missed some button presses. Experimental blocks lasted 9 s (with 3 trials per block) and fixation blocks lasted 14 s. Each run (consisting of 16 experimental blocks and 5 fixation blocks) lasted 214 s (3 min 34 s). Participants completed 2 runs.

#### Language network localizer experimental materials

##### Standard language localizer materials

The materials consisted of five sets, each set comprising 48 sentences and 48 nonword sequences, for a total of 240 sentences and 240 nonword sequences. The sentences were drawn from the Brown corpus (Bird & Looper, 2004; Francis & Kucera, 1964) and were selected to include a variety of syntactic constructions and topics. The nonwords were created using the ‘Wuggy’ software (https://github.com/WuggyCode/wuggy; the default parameters were used) so as to respect the phonotactic constraints of English. In cases where Wuggy was unable to generate a nonword candidate, we relied on one of the following strategies: i) broke down the word into composite words (for compound words) or morphemes, matched each composite word/morpheme to a nonword, and then reassembled those; ii) used one of the nonwords created for another word; or iii) created an English-sounding nonword ourselves. Any given participant saw one set of materials.

##### Speeded language localizer materials

The first 11 participants were presented with the materials from the standard version (ensuring that a different set was used). Approximately halfway through data collection, we created a new set of materials for the speeded language localizer in order to: i) generalize the findings to a new set of materials, and ii) avoid potential material overlaps between the standard and speeded localizer materials in future experiments. Hence, for the remaining 13 participants, we created five new sets each consisting of 48 sentences and 48 nonword sequences, for a total of 240 new sentences and 240 new nonword sequences. The sentences were again selected from the Brown corpus (Bird & Looper, 2004). In particular, we sampled 1,000 12 word-long sentences and then selected a set of 240 sentences that were not already included in the original set of materials, were syntactically and semantically diverse, and did not contain offensive/inappropriate content. The nonword strings were created as in the standard version.

##### Multiple Demand network localizer task

In addition to the language tasks, we included a non-linguistic demanding task: a spatial working memory task. The goal was two-fold. First, including a non-linguistic task allowed us to evaluate the *selectivity* of the language fROIs–defined by two versions of the localizer—for language processing (Fedorenko et al., 2011, 2024). And second, this task allowed us to examine brain responses to the conditions of the language localizer tasks in *another set of functional areas*: areas that comprise the domain-general Multiple Demand (MD) network (Duncan, 2010; Duncan et al., 2012; Fedorenko et al., 2013). This network supports executive functions like working memory and cognitive control. The spatial WM task has been previously established to robustly identify these areas at the individual-participant level (e.g., Blank et al., 2014; Mineroff, Blank et al., 2018; Shashidhara et al., 2020; Assem et al., 2020a; Malik-Moraleda, Ayyash et al., 2022). Although the areas of the MD network have been shown to not support any ‘core’ linguistic computations—like those related to lexical access, syntactic structure building, or semantic composition (e.g., Blank & Fedorenko, 2017; Diachek, Blank, Siegelman et al., 2020; Quillen et al., 2021; Shain, Blank et al., 2020, Shain et al., 2022)—their engagement has been reported for some cases of effortful perception and comprehension (e.g., Mattys & Wiget, 2011; MacGregor et al., 2022; Liu et al., 2022; see <u>Discussion</u>). We therefore wanted to evaluate the MD areas’ responses to speeded comprehension, to see whether this kind of processing difficulty draws on domain-general resources.

The spatial working memory task contrasted a *hard* condition with an *easy* condition in a standard blocked design with a counterbalanced condition order across runs (e.g., Fedorenko et al., 2011, 2013; Blank et al., 2014). On each trial (duration = 8 s), participants saw a fixation cross for 500 ms, followed by a 3×4 grid within which randomly generated locations were sequentially flashed (1s per flash) two at a time for a total of eight locations (*hard* condition) or one at a time for a total of four locations (*easy* condition). Then, participants indicated their memory for these locations in a two-alternative forced-choice paradigm via a button press (the choices were presented for 1,000 ms, and participants had up to 3 s to respond). Feedback, in the form of a green checkmark (correct responses) or a red cross (incorrect responses), was provided for 250 ms, with fixation presented for the remainder of the trial. Experimental blocks lasted 32 s (with 4 trials per block) and fixation blocks lasted 16 s. Each run (consisting of 12 experimental blocks and 4 fixation blocks) lasted 448 s (7 min 28 s). Participants completed 2 runs.

23 of the 24 participants completed the MD localizer in the same scanning session as the standard language localizer; the remaining participant—in a separate session (98 days apart).

### fMRI data acquisition, preprocessing and first-level analysis

#### fMRI data acquisition

Structural and functional data were collected on the whole-body, 3 Tesla, Siemens Trio scanner 32-channel head coil, at the Athinoula A. Martinos Imaging Center at the McGovern Institute for Brain Research at MIT. T1-weighted, Magnetization Prepared RApid Gradient Echo (MP-RAGE) structural images were collected in 176 sagittal slices with 1 mm isotropic voxels (TR = 2,530 ms, TE = 3.48 ms, TI = 1100 ms, flip = 8 degrees). Functional, blood oxygenation level dependent (BOLD) data were acquired using one of three similar sequences (denoted as sequence A, B, C). The data from the majority of participants (22 out of 24) were acquired using sequence A which we describe in this paragraph. See specifications of sequences B and C in SI Table 1 (importantly, scanning sequence is kept constant in all comparisons between the standard and the speeded versions of the localizer besides in a single participant). Sequence A was an SMS EPI sequence (with a 90 degree flip angle and using a slice acceleration factor of 2), with the following acquisition parameters: fifty-two 2 mm thick near-axial slices acquired in the interleaved order (with 10% distance factor), 2 mm × 2 mm in-plane resolution, FoV in the phase encoding (A ≫ P) direction 208 mm and matrix size 104 × 104, TR = 2,000 ms and TE = 30 ms, and partial Fourier of 7/8. The first 10 s of each run were excluded to allow for steady state magnetization.

#### fMRI preprocessing

fMRI data were analyzed using SPM12 (release 7487), CONN EvLab module (release 19b; https://web.conn-toolbox.org/resources/conn-extensions/evlab), and custom MATLAB scripts. Each participant’s functional and structural data were converted from DICOM to NIfTI format. All functional scans were coregistered and resampled using B-spline interpolation to the first scan of the first session (Friston et al., 1995). Potential outlier scans were identified from the resulting subject-motion estimates as well as from BOLD signal indicators using default thresholds in CONN preprocessing pipeline (5 standard deviations above the mean in global BOLD signal change, or framewise displacement values above 0.9 mm; (Nieto-Castanon, 2020). Functional and structural data were independently normalized into a common space (the Montreal Neurological Institute [MNI] template; IXI549Space) using SPM12 unified segmentation and normalization procedure (Ashburner & Friston, 2005) with a reference functional image computed as the mean functional data after realignment across all timepoints omitting outlier scans. The output data were resampled to a common bounding box between MNI-space coordinates (−90, −126, −72) and (90, 90, 108), using 2 mm isotropic voxels and 4th order spline interpolation for the functional data, and 1 mm isotropic voxels and trilinear interpolation for the structural data. Last, the functional data were smoothed spatially using spatial convolution with a 4 mm FWHM Gaussian kernel.

#### First-level analysis

Effects were estimated using a General Linear Model (GLM) in which each experimental condition was modeled with a boxcar function convolved with the canonical hemodynamic response function (HRF) (fixation was modeled implicitly, such that all timepoints that did not correspond to one of the conditions were assumed to correspond to a fixation period). Temporal autocorrelations in the BOLD signal timeseries were accounted for by a combination of high-pass filtering with a 128 s cutoff, and whitening using an AR(0.2) model (first-order autoregressive model linearized around the coefficient a = 0.2) to approximate the observed covariance of the functional data in the context of Restricted Maximum Likelihood estimation (ReML). In addition to experimental condition effects, the GLM design included first-order temporal derivatives for each condition (included to model variability in the HRF delays), as well as nuisance regressors to control for the effect of slow linear drifts, subject-motion parameters, and potential outlier scans on the BOLD signal.

### Definition of functional regions of interest (fROIs)

Language and Multiple Demand (MD) fROIs were defined using a group-constrained subject-specific (GSS) approach (Fedorenko et al., 2010) where a set of spatial masks, or parcels, is combined with each individual subject’s localizer activation map, to constrain the definition of individual fROIs. The parcels delineate the expected gross locations of activations for a given contrast and are sufficiently large to encompass the variability in the locations of individual activations. Within each parcel, we selected the top 10% most localizer-responsive voxels, based on t-values.

To define the language fROIs, we used a set of five parcels derived from a group-level probabilistic activation overlap map for the *sentences* > *nonwords* contrast in 220 independent participants. The parcels included two regions in the left inferior frontal gyrus (LIFG, LIFGorb), one in the left middle frontal gyrus (LMFG), and two in the left temporal lobe (LAntTemp and LPostTemp). Following prior work (e.g., Blank et al., 2014), to define the right-hemisphere RH fROIs, the LH parcels were transposed onto the RH, but the individual LH and RH fROIs were allowed to differ in their precise locations within the homotopic parcels. Although the mask contained six parcels, we decided to exclude the left angular gyrus (LAngG) due to accumulating evidence that LAngG differs functionally from the rest of the core temporal and frontal areas and exhibits lower functional connectivity with those core areas (for discussion, see Shain, Chen, Paunov et al., 2022). To define the MD fROIs, we used a set of 20 parcels (10 in each hemisphere) derived from a group-level probabilistic activation overlap map for the *hard* > *easy* spatial working memory contrast (Fedorenko et al., 2013) in 197 independent participants. The parcels included symmetrical regions in frontal and parietal lobes, as well as a region in the anterior cingulate cortex. All parcels are available for download from https://evlab.mit.edu/funcloc/.

### Extraction of fMRI BOLD responses

We evaluated language and MD networks’ responses by estimating response magnitudes to the conditions of the standard and speeded language localizers in the individually defined fROIs. For each fROI in each participant, we averaged the responses across voxels to get a single value per participant per fROI per condition (i.e., the *sentences* and *nonwords* conditions for the language localizer tasks, and the *hard* and *easy* conditions for the MD localizer task). The responses to the conditions used to localize the areas of interest (e.g., the responses to the *sentences* and *nonwords* conditions in the language fROIs) were estimated using an across-runs cross-validation procedure, where one run of the standard or speeded language localizer was used to define the fROI and the other run of the same localizer version was used to estimate the response magnitudes. The procedure was repeated for each of the two run partitions, once where the first run was used for fROI definition and the second run was used for response estimation and once where the second run was used for fROI definition and the first run was used for response estimation. Finally, the estimates were averaged to derive a single value per participant per fROI per condition.

### Statistical analysis

All statistical analyses were performed in RStudio (version 2021.09.2) or Python (version 3.9.2). Linear mixed effects (LME) models were used to test for differences in BOLD response magnitudes and spatial correlations across localizer versions and conditions, while accounting for random effects from different participants and fROIs.

All LMEs reported in the paper were implemented in R using the *lmer* function from the **lme4** package (Bates et al., 2015; version 1.1-31). Fixed effects included the condition (*sentences* vs. *nonwords*) and the localizer version (standard vs. speeded). Their interaction was also included as a fixed effect where specified in order to test whether the effect of condition was different between the two localizer versions. Random intercepts were fit for participants and fROIs. Statistical significance testing was performed using the **lmerTest** package (Kuznetsova et al., 2017; version 3.1-3). Using this package, the t-statistic and associated p-values based on Satterthwaite’s approximation were computed. In cases where the interaction term was included, we compared the model with and without the *condition:version* interaction. Model comparison was performed using a likelihood ratio test implemented via the *anova* function in **lme4**, which yielded a *χ*^2^ test statistic.

For analyses involving a single dependent measure per condition (i.e. when comparing spatial correlations between activation maps across the two localizer versions), the fixed-effect included only the localizer version and random intercepts similarly included the participants and fROIs (e.g. *SpCorr LH language ∼ version + (1|participant) + (1|fROI)*). Statistical significance was again computed using **lmerTest** based on the t-statistics when comparing model parameters.

For each LME reported, we provide tables containing the model formulae, fixed effects regression coefficients (ꞵ-values), t-values, p-values, and random effects coefficients (SI 2E, 3D, 3E, 3F, 4B). R-squared values were computed using the *r.squaredGLMM* function from the **MuMIn** package (version 1.47.1).

In addition to LME analyses, pairwise comparisons between two conditions were conducted using two-sided paired or one-sample t-tests, as appropriate. These tests were implemented with the scipy.stats module in Python. All tests were two-tailed and statistical significance was assessed at *α* = 0.05.

## Results

We compared the fMRI BOLD responses from two versions of a language localizer task (a ‘standard language localizer’ and a ‘speeded language localizer’). The results are organized according to the following two questions: 1) Can speeded reading be used to reliably localize language-responsive areas in individual participants?, and 2) Does increased processing difficulty during speeded reading affect brain responses in the domain-general Multiple Demand brain network?

### 1. The speeded language localizer can reliably localize language-responsive areas in individual participants

#### 1.A. The activation topography is similar between the standard and speeded language localizer versions

Twenty-four participants completed a standard language localizer task (Fedorenko et al., 2010) and a speeded localizer task. In both tasks, they silently read sentences (the *sentences* condition) and sequences of nonwords (the *nonwords* condition) (see <u>Methods; fMRI tasks</u>).

The activation maps for the *sentences > nonwords* contrast are visually highly similar between the standard and speeded language localizers (**Figure 1A**; see SI 2D for corresponding fROI surface maps in the same sample participants). To quantify this similarity, we correlated voxel-wise activation patterns (restricted to the LH language parcels; see SI 2A for whole-brain correlations) across localizer runs and versions. The correlation values were Fisher-transformed and averaged across the five LH language parcels, leading to a single value for each comparison.

**Figure 1.**
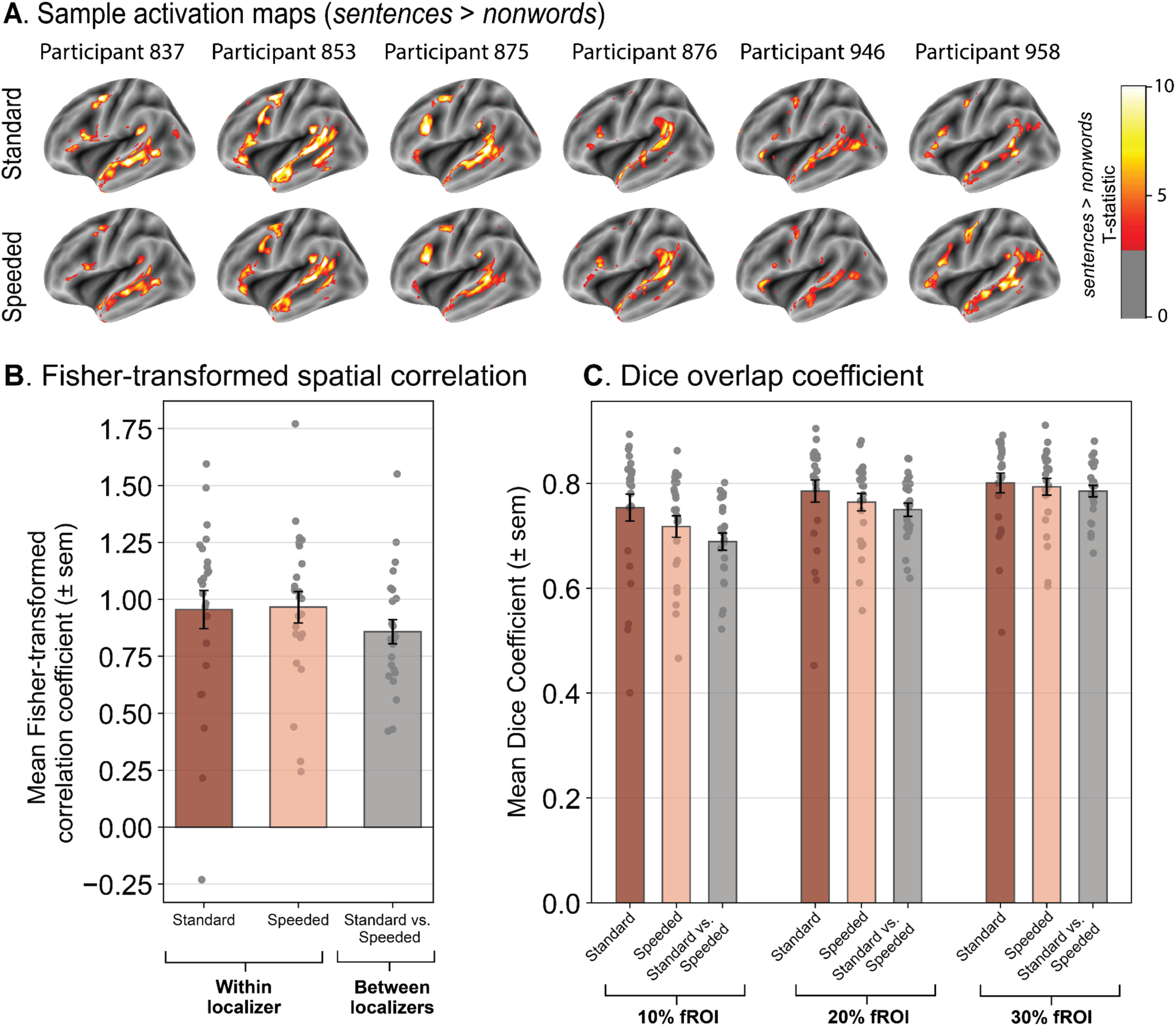
The activation topographies are highly similar between the standard and speeded language localizer versions. **(A)** Activation maps of the *sentences > nonwords* contrast in six sample participants for the standard language localizer (upper row) and the speeded language localizer (lower row). Activation maps are shown on the surface-inflated fsaverage template brain. The participant identifiers are numbers in the lab internal database and can be cross-referenced with the data tables on OSF. **(B)** The correlation of the voxel-wise activation patterns for the *sentences > nonwords* contrast within left-hemisphere (LH) language parcels (see <u>Methods; Definition of fROIs</u>) within and between localizer versions. The within localizer comparisons were performed by correlating the activation patterns between the two runs of the same localizer version (dark red bar = standard version; light red bar = speeded version; the data are averaged across participant and across five LH parcels within each participant), and the between localizer comparisons were performed by correlating the four pairwise combinations of runs between the two localizers (given two runs of each localizer; gray bar) and averaging them to obtain a single value. The dots correspond to the correlation values from individual participants (*n*=24). Error bars show the standard error of the mean across participants. **(C)** The Dice overlap coefficient for the *sentences > nonwords* contrast computed between different runs of the same localizer version or between different runs across the two localizer versions. The fROIs were defined as the top 10%, 20%, or 30% of the most language-responsive voxels in the five LH language parcels. As in panel B, the two red bars show the Dice coefficients within a localizer version (between two runs of the same localizer; the data are averaged across participants and across five LH parcels within each participant) and the gray bar shows the Dice coefficient between localizer versions (averaging across four pairwise between-run comparisons). The dots correspond to the coefficient values from individual participants (*n*=24). Error bars show the standard error of the mean across participants.

First, we correlated the activation patterns across the two runs *within* each localizer version. These values characterize the stability of the activation patterns for each version and also delimit the similarity that could be obtained between the two localizer versions. The within-localizer Fisher-transformed correlations were high for both versions: 0.955 and 0.966 for the standard and speeded versions, respectively (**Figure 1B**; left bars), and did not statistically differ from each other (*speeded* > *standard*; ꞵ=0.011, t=0.226, p=0.821 via linear mixed effects (LME) modeling). Next and critically, we correlated the activation patterns between the two versions of the localizer. To match the amount of data to the within-version comparisons, we correlated activations for each run of the standard version with each run of the speeded version (four pairwise combinations, given two runs of each localizer version; **Figure 1B**; the two *within* bars). The between-localizer Fisher-transformed correlation coefficient was 0.859 (**Figure 1B**; the *between* bar: the average of the four pairwise combinations of runs between the standard and speeded versions). To statistically compare the within- vs. between-version correlations, we modeled the average within-version and between-version correlation coefficients in an LME model with a fixed effect for comparison type (within vs. between), and random intercepts for participants and parcels. The similarity of the activations *within* a given localizer version was statistically higher than between localizer versions **(Figure 1B)**, with a relatively small effect size (**Figure 1B**; *within > between;* ꞵ=0.102, t=2.763, p=0.006), in line with both within and between correlations being high.

In a complementary analysis, we quantified the extent of voxel overlap between functional regions of interest (fROIs) using the Dice coefficient (Dice, 1945). The results mirrored the spatial correlation analyses above. The overlap between the *sentences > nonwords* fROIs, defined as the top 10% of language-responsive voxels, was high for both within and between comparisons (0.754 and 0.718 for the within comparisons for the standard and speeded versions, respectively; and 0.689 for the between comparison), but slightly higher across the runs within a localizer version than between localizer versions (ꞵ=0.047, t=3.291, p=0.001) (**Figure 1C**). For fROIs of larger size (e.g., fROIs defined as the top 20% or 30% of most language-responsive voxels within the parcels), the within vs. between differences get smaller (20%: ꞵ=0.025, t=2.281, p=0.023; 30%: ꞵ=0.012, t=1.238, p=0.217), which suggests that although the peaks of the activation topographies are slightly more similar within a localizer version than between the two versions, the overall topographies are highly similar (see SI 2B for Dice coefficient comparisons across a larger range of fROI thresholds).

#### 1.B. The fROIs defined by the speeded language localizer respond at least as strongly and as selectively during language processing as the fROIs defined by the standard localizer

Having established that the activation topographies are similar across the localizer versions (**Figure 1**), we examined the magnitude of the BOLD responses for the *sentences* and *nonwords* conditions across the two versions in the fROIs defined by the standard approach of selecting top 10% of most language-responsive voxels within five broad, anatomical parcels (**Figure 2A**; see <u>Methods; Extraction of fMRI BOLD responses</u>). **Figure 2B** shows the average BOLD responses across the five LH language fROIs and **Figure 2C** shows the responses for each of the five fROIs individually. To statistically compare the two localizers, the BOLD responses were modeled in an LME with fixed effects for condition (*sentences* vs. *nonwords*) and localizer version (standard vs. speeded) and random intercepts for participants and fROIs.

**Figure 2.**
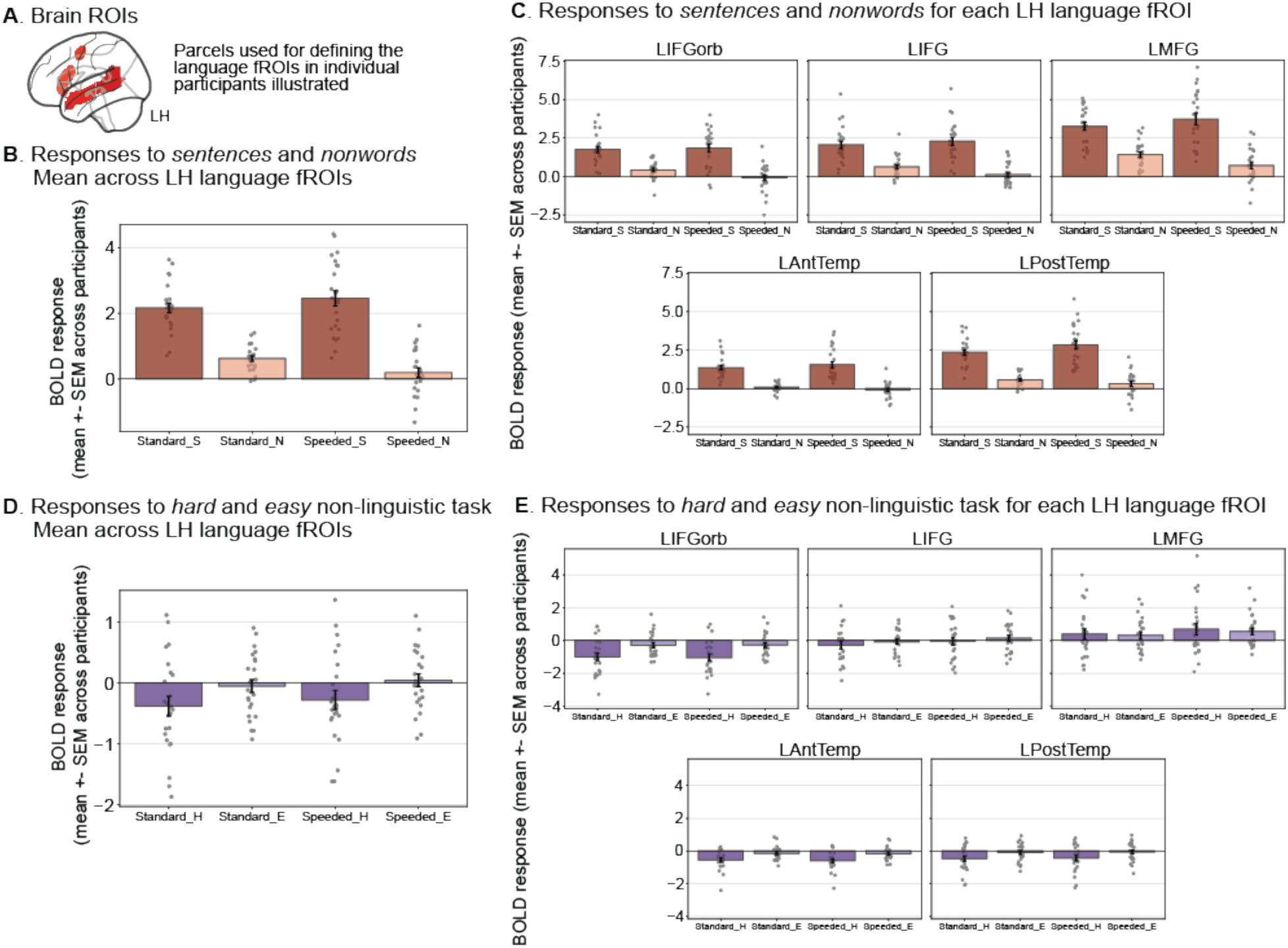
The speeded language localizer elicits a greater *sentences* > *nonwords* effect than the standard language localizer and the fROIs defined by the speeded localizer are similarly selective for language relative to a demanding non-linguistic task. **(A)** To define language fROIs, we used a set of masks (‘language parcels’; shown on the volumetric MNI152 template brain where all analyses were performed) within which most or all individuals in prior studies showed activity for the language localizer contrast in large samples (e.g., Fedorenko et al., 2010; Lipkin et al., 2022). We defined the LH language fROIs as the most language-responsive voxels (top 10%) within the borders of these five parcels for each participant and measured the BOLD response magnitude in these fROIs in a cross-validated manner (see <u>Methods; Definition of fROIs</u>). **(B)** Mean BOLD response to the language localizer conditions (S=*sentences*, N=*nonwords*) for the standard and speeded localizer versions averaged across the five LH language fROIs. **(C)** Mean BOLD response to the localizer conditions for each LH language fROI. **(D)** Mean BOLD response to a spatial working memory task consisting of two conditions, a *hard* condition (H) and an *easy* condition (E), averaged across the five LH language fROIs defined by the speeded and standard language localizers. See SI 3A for evidence that the spatial working memory task elicited a robust *hard* > *easy* response in the MD network fROIs. **(E)** Mean BOLD response to the *hard* and *easy* spatial working memory task conditions for each LH language fROI. In all panels, dots correspond to the responses of individual participants. Error bars show the standard error of the mean across participants. See SI 3E for responses in the right-hemisphere (RH) language fROIs (which also show a reliable *sentences > nonwords* effect, similar to the LH fROIs, although the responses are overall weaker).

As expected, the effect of condition (estimated in independent data) was highly significant (*sentences > nonwords*, ꞵ=1.897, t=23.570, p<0.0001) (**Figure 2B,C**); in contrast, the main effect of localizer version was not significant (*speeded* > *standard*, ꞵ=-0.068, t=-0.851, p=0.395). To further examine whether the standard and speeded versions differed with respect to their responses to *sentences* and *nonwords*, we used a similar LME as above but also included an interaction term between condition and localizer version. We tested for significance of the interaction using a likelihood-ratio test with a Chi Square test statistic (*χ*^2^). Indeed, the interaction was significant (*χ*^2^=20.653, p<0.0001), suggesting that the responses to the two conditions (*sentences* and *nonwords*) differed between localizer versions. To better understand this difference, we examined the size of the *sentences* > *nonwords* contrast and found that it was greater in the speeded localizer compared to the standard localizer (*speeded > standard*; ꞵ=0.723, t=7.817, p<0.0001). As can be seen in **Figure 2B,C**, the response to the *sentences* condition was higher in the speeded localizer compared to the standard localizer (*speeded > standard*; ꞵ=0.293, t=2.407, p=0.017), and the response to the *nonwords* condition was lower in the speeded localizer compared to the standard localizer (*speeded > standard*; ꞵ=-0.430, t=-5.106, p<0.0001). Taken together, these analyses show that the fROIs defined by the speeded language localizer show a larger *sentences* > *nonwords* effect compared to the standard localizer, due to both higher responses to *sentences* and lower responses to *nonwords* in the speeded version.

Because of the increasing interest in the field of language research in non-canonical language regions (e.g., Li et al., 2024; Wang et al., 2025; Tuckute et al., 2025), we also examined responses in the so-called *extended language network*, encompassing regions on the ventral temporal surface, in the medial frontal cortex, and in the cerebellum; Wolna et al., 2025). We found that the speeded localizer successfully identifies the regions of the extended language network and produces a similar response profile to the standard localizer (**SI 3G**), demonstrating that the speeded version can be used to identify not only the core language areas but also non-canonical language-responsive areas. Following a reviewer’s request, we additionally examined responses to language in the Default Mode Network (DMN), which has been implicated in semantic processing (e.g., see Fernandino & Binder, 2024; cf. Humphreys et al., 2015). Using the same individual fROI approach as in our analyses of the language areas and using the *easy > hard* contrast from the spatial working memory task to define the DMN regions (following Mineroff, Blank et al., 2018), we did not find substantial responses to language in the DMN regions for either localizer version (with the exception of the left anterior temporal DMN region, which overlaps with the anterior temporal language parcel) (**SI 3H**).

Next, we examined the selectivity of the language fROIs defined by both the standard and the speeded localizers for language processing relative to a non-linguistic demanding cognitive task. Prior work has established that language-responsive brain areas (as defined by standard versions of the language localizer) are highly selective for language relative to diverse non-linguistic inputs and tasks (e.g., Fedorenko et al., 2011; Ivanova et al., 2020, 2021; Chen et al., 2023; for reviews, see Fedorenko & Blank, 2020; Fedorenko et al., 2024). Here, we investigated whether the fROIs defined by the speeded language localizer exhibit a similar degree of selectivity. To do so, we collected brain responses during a spatial working memory task (see <u>Methods; fMRI tasks</u>) and examined BOLD response magnitudes to the *hard* and *easy* conditions in the LH language regions, defined by the standard versus speeded language localizers (**Figure 2D,E**). As expected given the high topographic overlap between the two localizer versions reported in Section 1, both sets of fROIs showed selectivity for language, with no response during the cognitively demanding spatial working memory task (standard localizer: *hard*: t=-2.376, p=0.026; *easy*: t=-0.505, p=0.618 via two-sided, one-sample t-test against zero; speeded localizer: *hard*: t=-1.803, p=0.085; *easy*: t=0.397, p=0.694). This lack of response in the language areas is in sharp contrast with the Multiple Demand areas, which respond strongly to both conditions, and show a clear *hard* > *easy* effect (SI 3A).

Finally, in addition to the analyses reported in 1.A and 1.B above, we tested whether the BOLD response magnitudes from the fROIs defined by the standard versus speeded localizers were stable over time (across runs (SI 3B) and—for two participants who completed the localizers several times—across scanning sessions (SI 3C)). This is important to know given that BOLD response magnitudes are often used in individual-differences investigations that aim to relate neural measures to behavior (e.g., Mahowald & Fedorenko, 2016; Assem et al., 2020a; Kong et al., 2020). We found that the magnitudes were indeed highly stable within participants over time.

### 2. Speeded sentence reading engages the domain-general Multiple Demand (MD) system to a greater extent than standard reading

In addition to examining responses in the language network (Section 1.B), we investigated responses in the domain-general Multiple Demand (MD) network. This network supports computations related to goal-directed behaviors and is recruited during a broad array of cognitively demanding tasks (e.g., Duncan, 2010; Duncan et al., 2012; Fedorenko et al., 2013; Shashidhara et al., 2019; Assem et al., 2020b; Duncan et al., 2020). Of most relevance to the current investigation, the MD network appears to be engaged in some cases of effortful comprehension, including processing speech in noisy conditions or following acoustic degradation (Mattys & Wiget, 2011; MacGregor et al., 2022; Liu et al., 2022), processing accented speech (Adank & Janse, 2010; Janse & Adank, 2012; Adank et al., 2012; Banks et al., 2015), processing languages that one is not fully proficient in (Malik-Moraleda, Jouravlev et al., 2024; Wolna et al., 2024), and processing linguistic inputs that are not syntactically well-formed (Kuperberg et al., 2003; Nieuwland et al., 2012; Mollica et al., 2020; Tuckute et al., 2024b; Kauf et al., 2024). However, understanding the full range of conditions under which the MD network is recruited during language processing remains an important research goal, and is critical for deciphering the nature of the MD network’s contributions to language.

Following prior work (e.g., Malik-Moraleda, Ayyash et al., 2022), we defined MD fROIs (10 in each hemisphere; **Figure 3A**) using the *hard > easy* contrast of the spatial working memory task described in the previous section (Section 1.B; and <u>Methods; fMRI tasks</u>). We then examined the responses to the *sentences* and *nonwords* conditions across the two versions of the language localizer to test whether speeded reading taxes the MD network. (For validation that the MD fROIs behave as expected, i.e., show a reliably greater response to the hard spatial working memory condition compared to the easy one, see SI 3A.)

**Figure 3.**
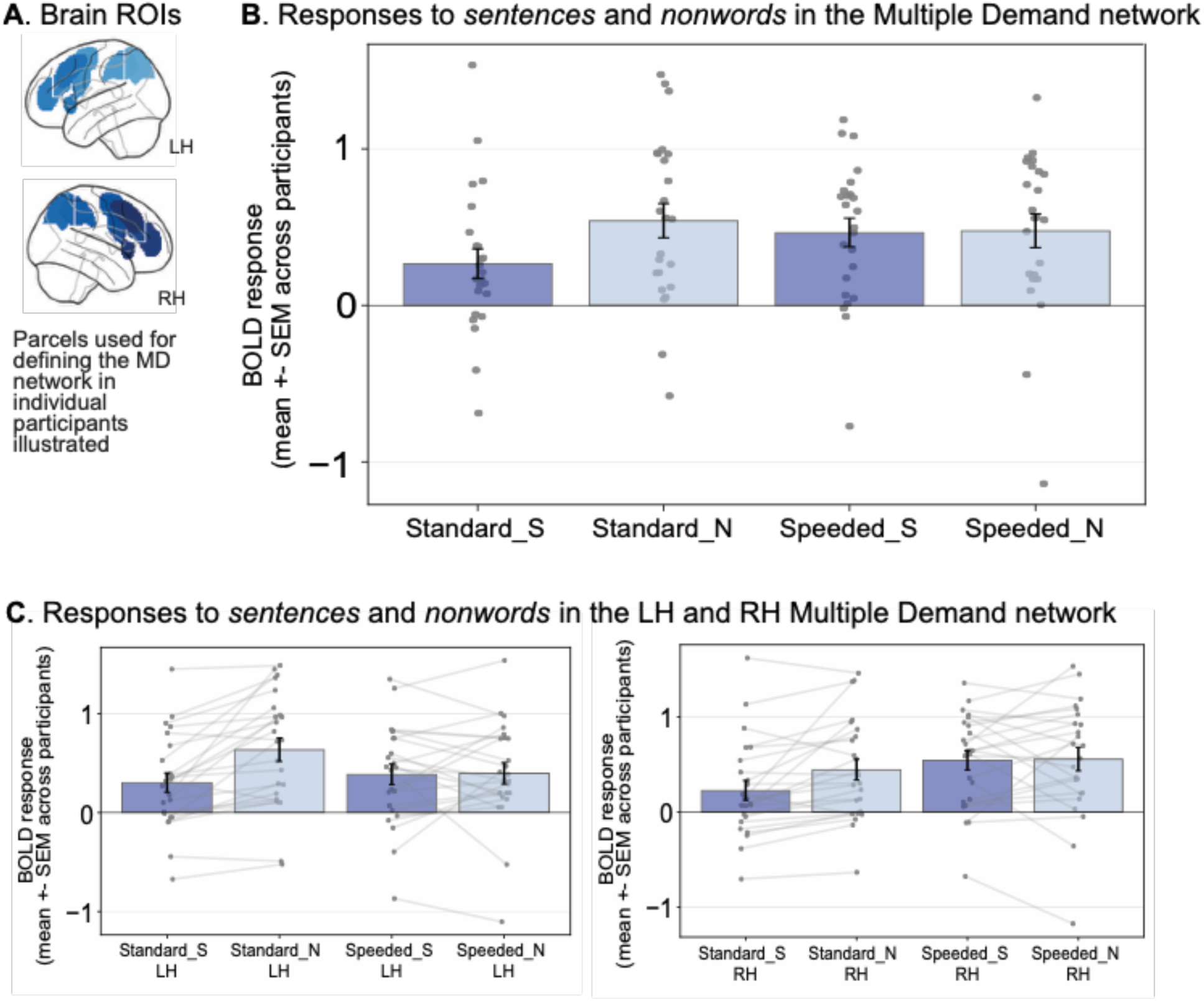
The Multiple Demand (MD) network is more engaged in speeded sentence reading compared to standard sentence reading. **(A)** To define MD fROIs, we used a set of masks (‘MD parcels’; shown on the volumetric MNI152 template brain) within which most or all individuals in prior studies showed activity for the MD *hard* > *easy* spatial working memory contrast in large samples (e.g., Diachek, Blank, Siegelman et al., 2020). We defined the LH and RH MD fROIs as the most working-memory-responsive voxels (top 10%) within the borders of the twenty parcels for each participant, and measured the BOLD response magnitude in these fROIs in a cross-validated manner (see <u>Methods; Definition of fROIs</u>). **(B)** Mean BOLD response to the language localizer conditions (S=*sentences*, N=*nonwords*) for the standard and speeded localizer versions averaged across the twenty LH/RH MD fROIs (see SI 4A for individual fROIs). **(C)** Mean BOLD response to the language localizer conditions (S=*sentences*, N=*nonwords*) for the standard and speeded localizer versions averaged across the LH and RH MD fROIs. Light gray lines connect the responses to the two conditions for a given participant and localizer version. In all panels, dots correspond to the responses of individual participants. Error bars show the standard error of the mean across participants.

The BOLD response magnitudes for the *sentences* and *nonwords* conditions across both localizer versions are shown in **Figure 3B** for the average of the ten left and right hemisphere MD fROIs and **Figure 3C** for each hemisphere separately (see SI 4A for each of the twenty fROIs individually). In line with prior work (e.g., Fedorenko et al., 2013; Diachek, Blank, Siegelman et al., 2020), the MD fROIs showed a robust *nonwords* > *sentences* effect in the standard language localizer (*sentences > nonwords,* ꞵ=-0.275, t=-7.547, p<0.0001). In contrast, in the speeded version, reading of *nonwords* did not engage the MD network to a greater extent than reading of *sentences* (*sentences* > *nonwords*, ꞵ=-0.012, t=-0.283, p=0.777). As evident from **Figure 3C**, some participants exhibited higher MD network engagement in the *nonwords* condition, whereas others exhibited the opposite pattern. To statistically compare the responses to the two localizers, the BOLD responses were modeled in an LME with fixed effects for condition (*sentences* vs. *nonwords*) and localizer version (standard vs. speeded) and random intercepts for participants and fROIs. Using likelihood ratio tests, we confirmed a significant interaction between condition and localizer version (*χ*^2^=18.274, p<0.0001), suggesting that the MD network was engaged differently by the two localizers. In particular, the MD network was more engaged in the *sentences* condition during the speeded localizer compared to the standard localizer (*speeded* > *standard*, ꞵ=0.200, t=4.584, p<0.0001), whereas the responses to the *nonwords* condition did not reliably differ between the two versions (*speeded > standard*, ꞵ=-0.064, t=-1.519, p=0.129). In summary, speeded sentence reading was more effortful than slower-paced reading, and under the speeded-reading conditions, no *nonwords > sentences* effect was observed.

## Discussion

In cognitive neuroscience, there is a growing recognition of inter-individual differences in the precise functional topographies, especially in the association cortex (e.g., Frost & Goebel, 2012; Tahmasebi et al., 2012; Braga et al., 2020; Gratton & Braga, 2021; DiNicola et al., 2023; Du et al., 2024). We here show that a standard localizer for the language network (Fedorenko et al., 2010) can be halved in time by using speeded reading, and that the speeded-reading-based contrast is even more robust than the one based on standard-paced reading. In the remainder of the Discussion, we elaborate on these findings and their implications.

### 1. Robustness and generalizability of the language localizer

The standard language localizer (Fedorenko et al., 2010) investigated in our study has been widely used over the past decade (e.g., Fedorenko et al., 2010; Mahowald & Fedorenko, 2016; Braga et al., 2020; Lipkin et al., 2022; Du et al., 2024). The localizer contrasts the reading of well-formed sentences versus sequences of nonwords. The brain areas identified by this contrast have been shown to be robust across different linguistic materials (e.g., Fedorenko et al., 2010)—an effect we replicate here—and tasks (e.g., Diachek, Blank, Siegelman et al., 2020; Gao et al., 2025). Moreover, this contrast generalizes well to the auditory and audio-visual presentation modalities (e.g., Fedorenko et al., 2010; Scott et al., 2017; Olson et al., 2023) and works well across typologically diverse languages (Richardson et al., 2020; Malik-Moraleda et al., 2022; Terhune-Cotter et al., 2023) and for diverse populations, including children (Hiersche et al., 2024; Ozernov-Palchik, O’Brien et al., in press), older healthy adults (Billot, Jhingan et al., 2025), and individuals with stroke aphasia (Billot, 2023; Clercq et al., 2024; Billot et al., in prep). In the current study, we show that the reading version of the localizer is robust to presentation speed, in line with past behavioral work showing the ability to understand language at fast speeds when presented word-by-word in a rapid serial visual presentation (RSVP) paradigm (e.g., Forster, 1970; Potter et al., 1980, 1986; Potter, 2012; Mollica & Piantadosi, 2017), and in line with prior group-averaging fMRI studies (Vagharchakian et al., 2012; Benjamin & Gaab, 2012). In the speeded version that we evaluated, each word was presented for 200 ms (compared to 450 ms in the standard localizer, i.e., ∼56% faster), and we demonstrate that language areas in individual participants can be reliably localized using this version.

### 2. The speeded language localizer shows at least as strong selectivity for language relative to the control condition and a non-linguistic demanding task

In the current work, we first established that the voxel-level activation topographies were highly similar between the standard and speeded language localizers, and then demonstrated that the response magnitudes in fROIs defined by each localizer version were also similar both in their responses to language and a control condition, and in their selectivity for language relative to a non-linguistic working memory task (e.g., Duncan, 2010; Fedorenko et al., 2013). Moreover, the speeded localizer is actually more effective than the standard version given that it better differentiates the critical language condition and the control condition. Specifically, the speeded localizer elicited a stronger response to the *sentences* condition, possibly due to an increase in attentional demands or processing difficulty (but see next discussion section), and a weaker response to the control condition (*nonwords*). The reduced response to nonwords may be due to the increased challenge of reading nonwords quickly which in turn might reduce the accessibility of information about their phonotactic properties (e.g., Regev et al., 2024). Thus, the speeded localizer produced a response profile with at least as strong responses to language as the standard localizer. Additionally, the areas identified by the speeded localizer were selective for language relative to a non-linguistic spatial working memory task, similar to the profile of the areas identified using the standard localizer (see Fedorenko & Blank, 2020 and Fedorenko et al., 2024 for reviews).

We also found that the size of the *sentences > nonwords* contrast was stable across runs for the speeded localizer version, similar to the standard version, which suggests that the speeded localizer can also be used in studies that relate neural markers to behavior or genetics to study individual differences (e.g., Mahowald & Fedorenko, 2016; Assem et al., 2020a; Kong et al., 2020).

### 3. Contributions of the Multiple Demand (MD) network to language comprehension

The Multiple Demand (MD) network is broadly implicated in cognitively demanding tasks and goal-directed action, showing strong responses to diverse executive function tasks (Duncan & Owen, 2000; Duncan, 2010; Duncan et al., 2012; Fedorenko et al., 2013; Shashidhara et al., 2019b; Assem et al., 2020b; Duncan et al., 2020) as well as during some domains of reasoning, like arithmetic reasoning (e.g., Monti et al., 2009; Fedorenko et al., 2013; Amalric & Dehaene, 2019) and understanding computer code (e.g., Ivanova et al., 2020; Liu et al., 2020). Further, language paradigms where comprehension/production is accompanied by extraneous task demands—e.g., performing meta-linguistic judgments or answering comprehension questions—have been shown to engage frontal and parietal areas whose topography is consistent with that of the MD network (e.g., Cutting et al., 2006; Christodoulou et al., 2014). Similar studies that have included an MD network localizer confirm that these task effects occur in the MD areas (e.g., Diachek, Blank, Siegelman et al., 2020; Gao et al., 2025). However, during naturalistic comprehension of even syntactically complex stimuli, the MD network is not engaged, at least when linguistic stimuli are presented auditorily, and the costs of language processing are localized to the language-selective system (Diachek, Blank, Siegelman et al., 2020; Quillen et al., 2021; Wehbe et al., 2021; Shain et al., 2022; see review, see Fedorenko & Shain, 2021). Unlike passive listening to language, passive sentence reading, even at typical speeds, sometimes elicits a slightly above-baseline response (e.g., Gao et al., 2025). We see such a response here for our standard version as well. This effect may have to do with the fact that unlike listening, reading is a later-acquired skill and may be associated with some amount of cognitive effort even for proficient readers.

In contrast to the costs associated with *linguistic* demands specifically (e.g., processing unexpected elements or building non-local inter-word dependencies; Shain, Blank et al., 2020; Shain et al., 2022), some cases of effortful comprehension, even without external task demands, appear to engage the MD network. Such cases include listening to speech in noisy conditions (Mattys & Wiget, 2011; MacGregor et al., 2022; Liu et al., 2022), processing accented speech (Adank & Janse, 2010; Janse & Adank, 2012; Adank et al., 2012; Banks et al., 2015), processing sentences in a language that one is not fully proficient in (Malik-Moraleda, Jouravlev et al., 2024; Wolna et al., 2024), and processing linguistic inputs that are not syntactically well-formed (Kuperberg et al., 2003; Nieuwland et al., 2012; Mollica et al., 2020; Tuckute et al., 2024b; Kauf et al., 2024). A possible generalization about these cases is that they all involve difficulty extracting a syntactically parsable word sequence from perceptual linguistic inputs.

Here, we present another case where passive language comprehension engages the MD network: speeded reading (for earlier evidence, see Vagharchakian et al., 2012, although the evidence is indirect as no independent MD localizer is included). The MD regions’ response during the *sentences* condition was ∼43% higher in the speeded version compared to the standard version (cf. a much smaller difference observed in the language regions: a ∼16% increase for the speeded version). Interestingly, in some previously reported cases, the linguistic condition that engages the MD network to a greater extent elicits a *lower* response in the language areas. For example, Malik-Moraleda, Jouravlev et al. (2024) show that comprehension of languages in which participants have low proficiency engages the MD network more strongly than higher-proficiency languages, but elicits a *lower* response in the language network. In contrast, the speeded sentence reading condition elicited a higher response compared to the normal-speed reading condition in both the MD network and the language network. This pattern may be taken to suggest that the generalization above—that the MD network gets engaged when it is difficult to extract a syntactically parsable word sequence from perceptual inputs—is not correct: this kind of difficulty should systematically lead to lower responses in the language network given that partially comprehensible stimuli should not be able to engage linguistic computations to the full extent (see Malik-Moraleda, Jouravlev et al., 2024 and Ozernov-Palchik, O’Brien et al., in press, for discussion). Thus, the precise contributions of the MD network during different kinds of effortful linguistic processing remain to be determined.

Finally, given the MD network’s stronger response during the speeded sentence reading condition but a similarly strong response during the nonword reading condition, the speeded localizer does not elicit a *nonwords > sentences* effect in the MD regions, in contrast to the standard language localizer (Fedorenko et al., 2013; Diachek, Blank, Siegelman et al., 2020). A practical implication is that it is not possible to use the *nonwords > sentences* contrast in the speeded version to localize the MD network (in addition to the language network) as is sometimes done (e.g., Shain, Blank et al., 2020). Whether the time saved by the speeded language localizer version is worth this trade-off of not being able to functionally define the MD regions using the same localizer will depend on the researcher’s goals.

### 4. Other efforts in cognitive neuroscience to develop efficient localizers

Functional localizers increase the sensitivity, functional resolution, and interpretability of research in cognitive neuroscience, but they take up precious time during the study. As a result, there is growing interest in making localizers more efficient. Two approaches have been taken to create more efficient localizers: i) reducing the number and/or the duration of experimental blocks, or ii) trying to optimize the critical stimuli so as to increase the size of the *critical > control* effect. Our approach falls into the first category: by increasing the speed of (visually) presenting linguistic materials (by ∼56%), we shortened experimental blocks from 18 s (3 6-second trials) to 9 s (3 3-second trials). (Note that although we retained the original 14 s fixation blocks for maximum comparability with the standard version, the fixation blocks could likely be shortened to 9 s, which would shave off another 30 s from the run’s duration.) Lee et al. (2024) also took the first approach, but instead of changing the presentation speed, they iteratively removed blocks and examined the consequences on brain responses. They showed that for a standard auditory language localizer based on the contrast of *intact speech > degraded speech* (as introduced in Scott et al., 2017) reducing the two-run protocol from ∼12 minutes (16 intact and 16 degraded blocks) to ∼6 minutes at 50% (16 blocks total) and even ∼3.5 minutes at 25% (8 blocks total) suffice for localizing the language regions.

The approach of stimulus optimization was used by Dodell-Feder et al. (2011), who analyzed responses to individual stimuli in a standard Theory of Mind (ToM) network localizer (Saxe & Kanwisher, 2003). Using a large dataset of a few hundred participants, they identified a) a subset of the critical-condition items (false belief stories) that elicit the highest response in the ToM brain areas, and b) a subset of the control-condition items (false photograph stories) that elicit the lowest response in the ToM areas. These subsets were used to create a highly efficient ToM localizer (see Chen, Kamps et al., 2024 for a related approach). Other studies have attempted to select stimuli that would be especially exciting for particular individuals based on their interests. For example, Olson, D’Mello et al. (2023) used language materials on topics of interest to different individuals with autism and found stronger responses in the language areas with those custom-selected stimuli. Finally, with the advent of neural networks that are predictive of brain responses (e.g., Yamins et al., 2014; Schrimpf et al., 2021, Tuckute et al., 2024a), it is now possible to create or select stimuli that elicit maximal responses in the target region/network (Bashivan et al., 2019; Xiao & Kreiman, 2020; Ratan Murty et al., 2021; Gu et al., 2023; Tuckute et al., 2024b). To our knowledge, these advances have not yet been leveraged in the creation of efficient localizers, but they certainly can and should be.

In addition to increasing the efficiency of a given localizer, another recent effort is to combine several localizers into a single experiment. For example, Marvi, Hutchinson et al. (2025) propose a multimodal localizer with simultaneous presentation of visual stimuli, such as faces, bodies, and scenes, which can be processed relatively bottom-up/automatically (e.g., Bugatus et al., 2017; Mur et al., 2025), and auditory stimuli, such as meaningless speech, sentences, and even short passages, which engage high-level ToM areas. This multimodal localizer is shown to elicit brain responses comparable to administering all the different contrasts in independent tasks.

Increasing localizer efficiency in all these ways is valuable given the increasing popularity of precision imaging approaches in cognitive neuroscience (Gordon et al., 2017; Naselaris et al., 2021; Gratton & Braga, 2021; Allen et al., 2022).

### 5. Limitations

A limitation of the speeded localizer is that it is dependent on the reading level of participants and the ability to read under speeded conditions. We took inspiration from prior behavioral evidence of successful linguistic processing under RSVP at presentation rates that are even faster than the 200ms/word used here (e.g., Forster, 1970; Potter et al., 1980, 1986; Potter, 2012; Mollica & Piantadosi, 2017). But of course, in those earlier studies, and in our study, the participants are proficient adult readers. The speeded localizer may be less suitable for children, older adults, and other populations that may have difficulties with reading at fast rates (e.g., Christodoulou et al., 2014).

A limitation of the current study design is that the standard language localizer was always presented prior to the speeded version (typically separated by other, unrelated tasks), so differences, or lack thereof, between the two versions could be confounded by task order. Although the experimental materials were always different between the versions, repeating a similar task within a session can modulate BOLD responses through practice with the task, practice-related changes in strategy, or reduced attention later in the session. However, these kinds of confounds should lead to lower BOLD responses later in the session (e.g., Landau et al., 2004; Grill-Spector et al., 2006). However, we observe that the response to the critical sentence condition in the speeded version is *higher* than the responses to sentences in the standard version despite being administered later in the session. It is quite likely that doing the reading in an RSVP paradigm at a slower rate makes it easier to perform it at a faster rate later, so performing the speeded version first may lead to an even stronger response to the sentence condition, but this possibility remains to be evaluated.

### 6. Conclusions

We hope that researchers working with populations who are proficient readers would benefit from this version of the language localizer, and that creating more efficient localizers, like this one, would lead to an even more widespread adoption of functional localization as a way to build a cumulative research enterprise where findings can be more straightforwardly compared across studies and labs.

## Data and code availability

The scripts for running the speeded language localizer as well as the associated analyses can be found here: https://github.com/el849/speeded_language_localizer/. The data can be found on OSF: https://osf.io/2vskh/.

## Author contributions

Conceptualization: G.T., E.F. Data curation: G.T., E.J.L., A.S. Investigation: G.T., E.J.L., A.S. Formal analysis: G.T., E.J.L. Visualization: G.T., E.J.L., Writing-original draft: G.T., E.J.L., E.F., Writing-review and editing: G.T., E.J.L., A.S., E.F.

## Funding and acknowledgements

We acknowledge the Athinoula A. Martinos Imaging Center at the McGovern Institute for Brain Research, MIT, including the technical team: Steve Shannon and Atsushi Takahashi. We also thank Agata Wolna for helpful discussions. GT was supported by the K. Lisa Yang ICoN Center Graduate Fellowship and the Chan Zuckerberg Initiative Foundation to establish the Kempner Institute at Harvard University. EF and this research were partially supported by funds from the McGovern Institute for Brain Research and the Simons Center for the Social Brain at MIT.

## Ethics

All participants gave informed written consent in accordance with the requirements of the MIT’s Committee on the Use of Humans as Experimental Subjects (COUHES).

## Supplementary Information

## SI 1: Details on fMRI acquisition sequences

**SI Table 1.** Functional MRI acquisition sequences. * One participant had the standard language localizer acquired using sequence B, and the speeded language localizer acquired using sequence A. For all remaining participants, the acquisition sequence was kept constant in all comparisons between the standard and the speeded versions of the language localizer. For acquisition of the Multiple Demand (MD) localizer task, all participants besides one, completed the MD localizer task in the same session as the language localizer tasks. The remaining participant completed the two language localizer tasks using sequence B, while the MD task was acquired using sequence A in a later session.

|  | **A** | **B** | **C** |
| --- | --- | --- | --- |
| **Number Participants** | 22* | 2* | 1 |
| **Sequence** |  |  |  |
| **Type** | SMS-EPI | EPI with GRAPPA | EPI with GRAPPA |
| **Flip Angle** | 90 | 90 | 90 |
| **Acceleration Factor** | 2 | 2 | 2 |
| **Acquisition Parameters** |  |  |  |
| **Number Slices** | 52 | 33 | 31 |
| **Slice Thickness** | 2mm | 4mm | 4mm |
| **In-plane Resolution** | 2mm x 2mm | 2.1mm x 2.1mm | 2.1mm x 2.1mm |
| **FoV** | 208mm | 200mm | 200mm |
| **FoV Matrix Size** | 104 x 104 | 96 x 96 | 96 x 96 |
| **TR** | 2000ms | 2000ms | 2000ms |
| **TE** | 30ms | 30ms | 30ms |

## SI 2: Information related to Results Section 1.A

### SI 2A: Whole-brain spatial correlation (supplementing language parcel correlations in Figure 1B and Figure 1C)

**SI Figure 2A:**
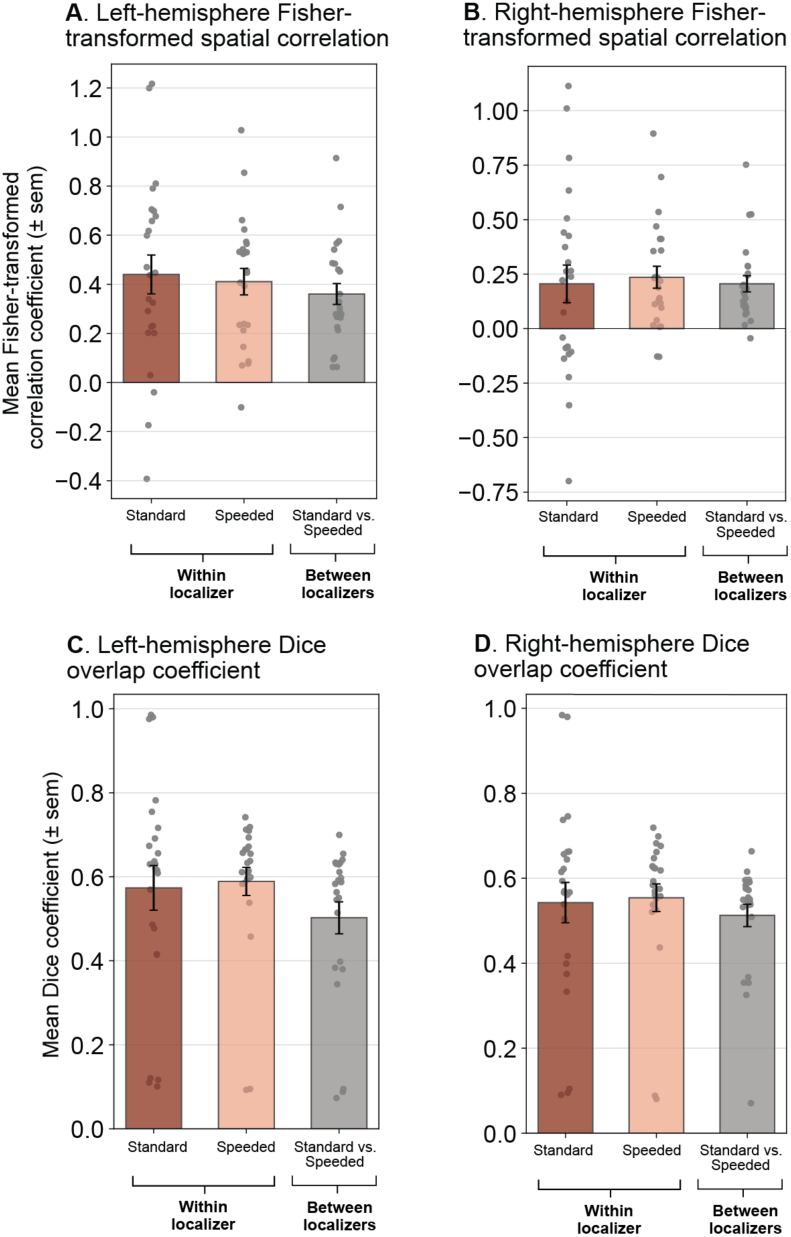
Correlation of whole-brain voxel-wise activation patterns and overlap coefficient within and between language localizer versions (for the left and right hemispheres separately). **(A-B)** We quantified the correlation of the voxel-wise activation patterns for the *sentences > nonwords* contrast within the whole left hemisphere (**panel A**) and right hemisphere (**panel B**) within localizer versions (between the two runs of the same localizer; red bars) and between localizer versions (for a total of four such pairwise combinations, given two runs of each localizer version; gray bar). In both panels, the bars show the average Fisher-transformed correlation coefficient across participants and individual points show the correlation values from individual participants (*n*=24). Error bars show the standard error of the mean across participants. **(C-D)** For an additional metric of the similarity of voxel-wise activation patterns for the *sentences* > *nonwords* contrast between the standard and speeded localizers, we computed the Dice coefficient overlap within the whole left hemisphere (**panel C**) and right hemisphere (**panel D**) within localizer versions (between the two runs of the same localizer; red bars) and between localizer versions (for a total of four such pairwise combinations, given two runs of each localizer version; gray bar). The Dice coefficient was computed as: 2 ∗ |Standard ∩ Speeded|/ (|Standard| + |Speeded|) for each hemisphere, where |Standard ∩ Speeded| denotes the number of voxels that were in the top 10% responsive voxels for both the standard and speeded localizer versions, |Standard| denotes the number of voxels in the top *10%* for the standard localizer version, and |Speeded| denotes the number of voxels in the top 10% for the speeded localizer version (Note: |Standard| = |Speeded| because the same parcels were used for both localizer versions). This computation provides a value between 0 and 1, where 0 indicates that the two localizer versions identified completely non-overlapping regions, and 1 indicates that the two localizer versions identified completely overlapping regions. In both panels, the bars show the average Dice coefficient across participants and individual points show the overlap values from individual participants (*n*=24). Error bars show the standard error of the mean across participants.

### SI 2B: Dice overlap coefficient between the standard and speeded language localizer versions

**SI Figure 2B.**
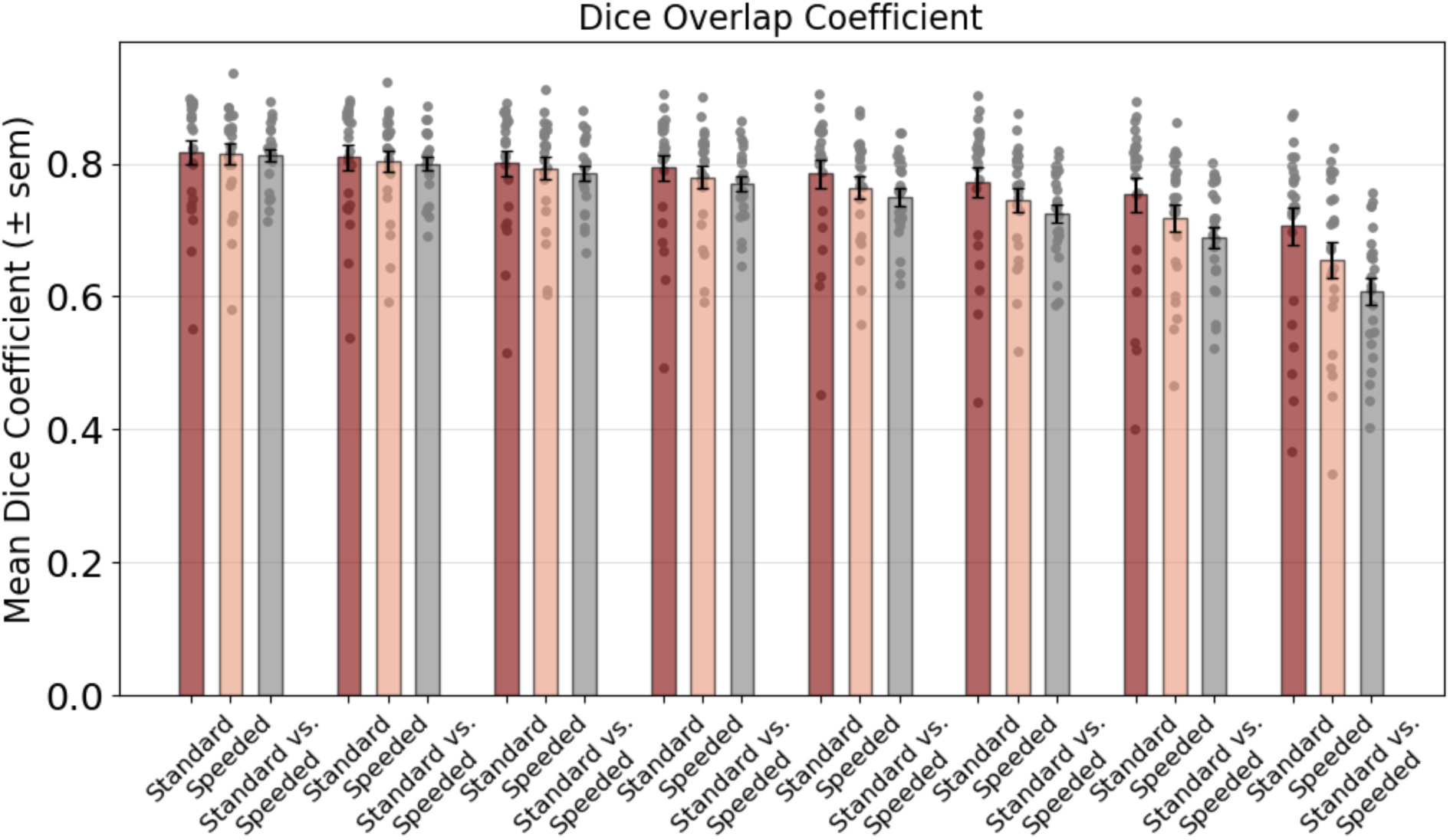
Dice coefficient overlap values across a range of fROI definition thresholds. We quantified the Dice coefficient overlap at a range of fROI definition thresholds. For each participant, the top *n*% most responsive voxels to the *sentences* > *nonwords* contrast in each of the LH language parcels were selected in both the standard and speeded localizer versions. *n* denotes the percentage threshold for fROI inclusion, and we show results for *n* = [5, 10, 15, 20, 25, 30, 35, 40]. The bars above show the average Dice coefficient across the five LH language fROIs. Note that in some participants (in particular for larger values of *n*), not all top *n*% voxels displayed a positive *sentences* > *nonwords* t-statistic. In this case, the voxels with negative t-statistic (i.e., opposite selectivity) were excluded from the Dice coefficient analyses. See the number of included voxels across the range of *n* in SI Table 2C.

**SI Table 2C.**
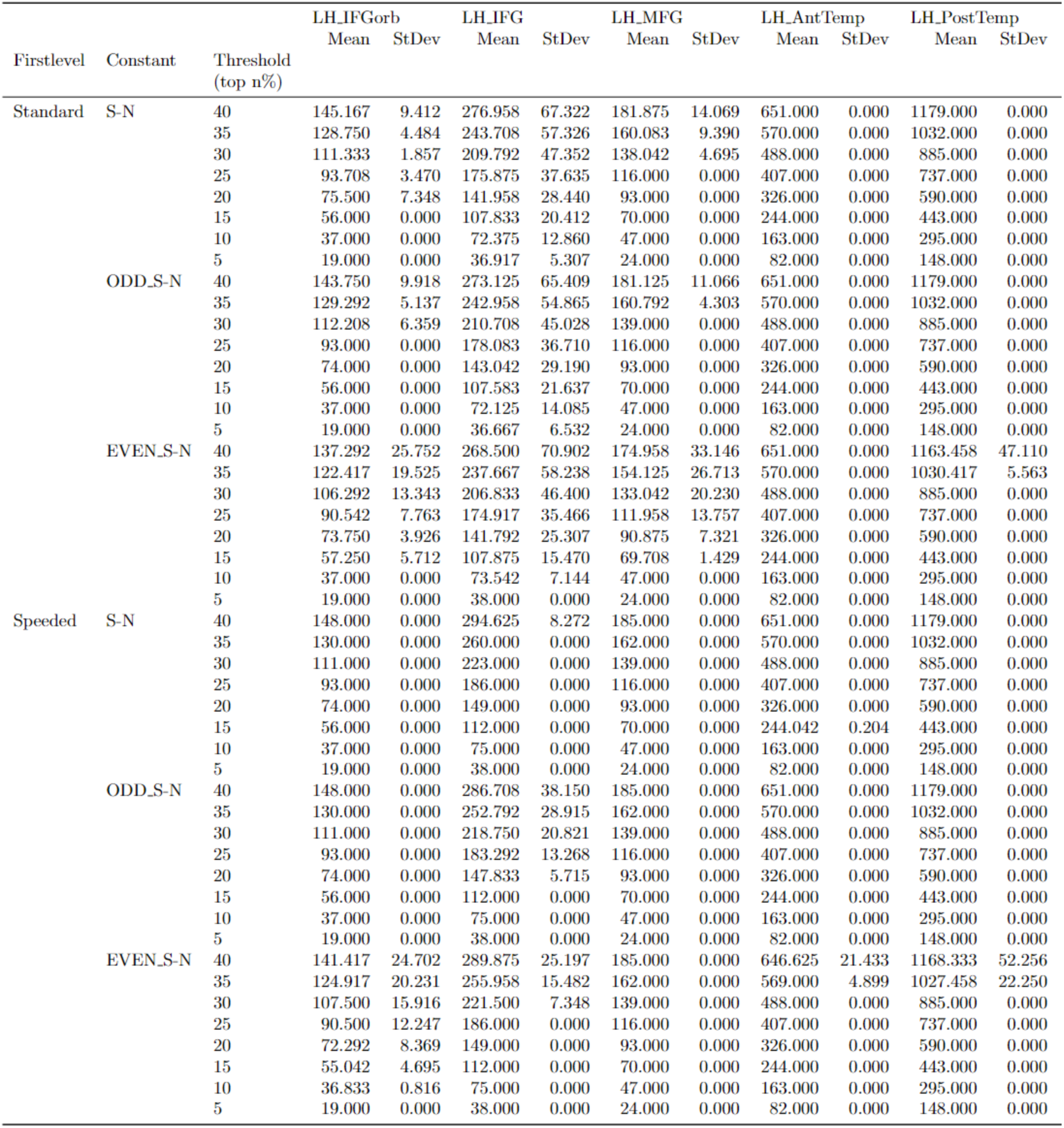
Number of included voxels across fROI definition thresholds for Dice overlap analyses. Mean and standard deviation for the number of voxels across participants in each left-hemisphere language fROI that were included in the fROIs for the Dice coefficient overlap analyses (i.e., voxels with a positive t-value corresponding to the *sentences > nonwords* contrast). For the Dice analyses, voxels that demonstrate the opposite selectivity (negative t-values) were excluded. The total number of voxels in the parcels were: LH_IFGorb: 370; LH_IFG: 743, LH_MFG: 462, LH_AntTemp: 1627, LH_PostTemp: 2948, and if no participants displayed negative t-values for the *sentences > nonwords* contrast the number of voxels included in the Dice analyses would always correspond to a given percentage threshold (*n*). As evidenced from the table, in most cases all *n* % voxels show positive t-values, but occasionally some voxels are excluded (in particular, for larger *n*).

### SI 2D: Sample participant fROI Maps

**SI Figure 2D.**
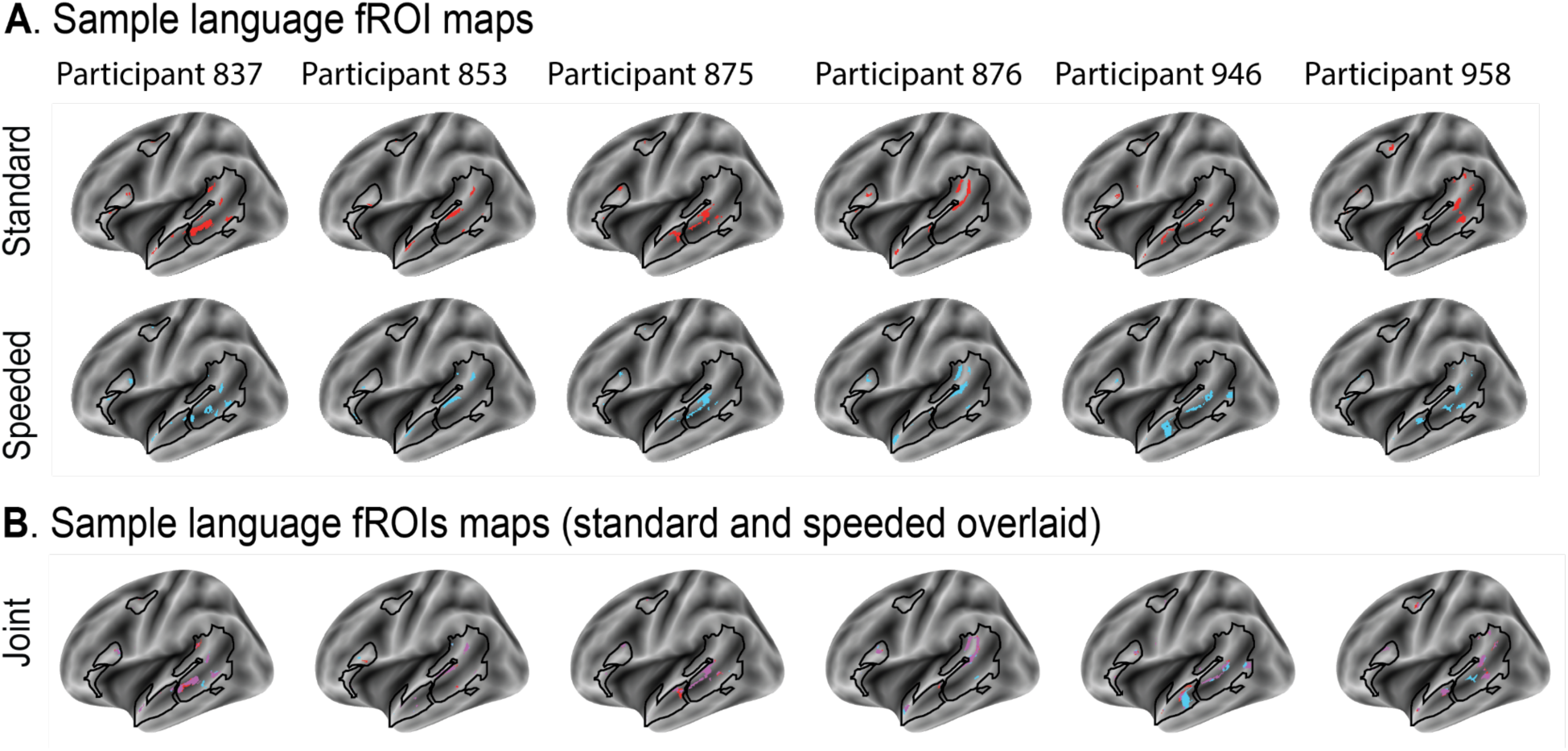
fROI maps for the standard and speeded language localizers. **(A)** Binary maps of the language fROIs in six sample participants for the standard language localizer (upper row in red) and the speeded language localizer (lower row in blue). fROIs were defined by selecting the top 10% most responsive voxels to the *sentences > nonwords* contrast (see <u>Methods; Definition of fROIs</u>). Maps are shown on the surface-inflated fsaverage template brain. The participant identifiers are numbers in the lab internal database and can be cross-referenced with the data tables on OSF. **(B)** Overlay of the language fROI maps from **(A)**. The fROI defined using the standard language localizer is shown in red, the fROI defined using the speeded language localizer is shown in blue. Regions that are overlapping in the two localizers are shown in purple.

### SI 2E: Statistics tables for Results Section 1.A

In the tables below, “SpCorr” denotes the Fisher-transformed spatial correlation coefficient. “within_between” denotes whether a spatial correlation coefficient was computed within localizer or between localizer versions. “participant” denotes each of the *n*=24 participants. “fROI” denotes each of the LH or RH regions of interest (five fROIs in each hemisphere).

|  | Estimate | Std. Error | df | t value | Pr(>\|t\|) |
| --- | --- | --- | --- | --- | --- |
| (Intercept) | 0.955 | 0.078 | 22.310 | 12.221 | 0.000 |
| VersionSpeeded | 0.011 | 0.047 | 211.788 | 0.226 | 0.821 |

$$SpCorr \text{ LH language} \sim within\_between + (1|participant) + (1|fROI)$$ Spcorr within localizer vs. between localizer versions, $r^2 = 0.432$
|  | Estimate | Std. Error | df | t value | Pr(>\|t\|) |
| --- | --- | --- | --- | --- | --- |
| (Intercept) | 0.859 | 0.074 | 24.566 | 11.601 | 0.000 |
| Conditionwithin | 0.102 | 0.037 | 331.871 | 2.763 | 0.006 |

*SpCorr language (LH wholebrain) ~ within\_between + (1|participant)* Spcorr within localizer vs. between localizer versions (left-hemisphere wholebrain), $r^2 = 0.5399$
|  | Estimate | Std. Error | df | t value | Pr(>\|t\|) |
| --- | --- | --- | --- | --- | --- |
| (Intercept) | 0.360 | 0.059 | 45.810 | 6.076 | 0.000 |
| Conditionwithin | 0.065 | 0.050 | 48.000 | 1.315 | 0.195 |

*SpCorr language (RH wholebrain) ~ within\_between + (1|participant)* Spcorr within localizer vs. between localizer versions (right-hemisphere wholebrain), $r^2 = 0.4235$
|  | Estimate | Std. Error | df | t value | Pr(>\|t\|) |
| --- | --- | --- | --- | --- | --- |
| (Intercept) | 0.206 | 0.060 | 53.012 | 3.406 | 0.001 |
| Conditionwithin | 0.015 | 0.056 | 48.000 | 0.265 | 0.792 |

*Dice coefficient ~ within\_between + (1|participant) + (1|fROI)* Dice within localizer vs. between localizer versions (top 10%), $r^2 = 0.344$
|  | Estimate | Std. Error | df | t value | Pr(>\|t\|) |
| --- | --- | --- | --- | --- | --- |
| (Intercept) | 0.689 | 0.028 | 13.617 | 24.527 | 0.000 |
| Conditionwithin | 0.047 | 0.014 | 331.866 | 3.291 | 0.001 |

Dice within localizer vs. between localizer versions (top 20%), $r^2 = 0.3726$
|  | Estimate | Std. Error | df | t value | Pr(>\|t\|) |
| --- | --- | --- | --- | --- | --- |
| (Intercept) | 0.750 | 0.023 | 13.771 | 32.806 | 0.000 |
| Conditionwithin | 0.025 | 0.011 | 331.882 | 2.281 | 0.023 |

Dice within localizer vs. between localizer versions (top 30%), $r^2 = 0.361$
|  | Estimate | Std. Error | df | t value | Pr(>\|t\|) |
| --- | --- | --- | --- | --- | --- |
| (Intercept) | 0.785 | 0.019 | 15.668 | 40.276 | 0.000 |
| Conditionwithin | 0.012 | 0.010 | 331.870 | 1.238 | 0.217 |

*Dice coefficient (LH wholebrain) ~ within\_between + (1|participant) + (1|fROI)* Dice within localizer vs. between localizer versions (left-hemisphere wholebrain top 10%), $r^2 = 0.0518$
|  | Estimate | Std. Error | df | t value | Pr(>\|t\|) |
| --- | --- | --- | --- | --- | --- |
| (Intercept) | 0.502 | 0.042 | 71.944 | 12.078 | 0.000 |
| Conditionwithin | 0.079 | 0.050 | 48.000 | 1.565 | 0.124 |

Dice within localizer vs. between localizer versions (left-hemisphere wholebrain top 20%), $r^2 = 0.3757$
|  | Estimate | Std. Error | df | t value | Pr(>\|t\|) |
| --- | --- | --- | --- | --- | --- |
| (Intercept) | 0.953 | 0.020 | 56.158 | 46.970 | 0.000 |
| Conditionwithin | -0.003 | 0.020 | 48.000 | -0.130 | 0.897 |

Dice within localizer vs. between localizer versions (left-hemisphere wholebrain top 30%), $r^2 = 0.2216$
|  | Estimate | Std. Error | df | t value | Pr(>\|t\|) |
| --- | --- | --- | --- | --- | --- |
| (Intercept) | 0.969 | 0.003 | 65.909 | 370.615 | 0.000 |
| Conditionwithin | -0.002 | 0.003 | 48.000 | -0.878 | 0.384 |

Dice coefficient (*RH wholebrain*) ∼ *within* *between* + (*1*|*participant*) + (*1*|*fROI*) Dice within localizer vs. between localizer versions (right-hemisphere wholebrain top 10%), $r^2 = 0.1371$
|  | Estimate | Std. Error | df | t value | Pr(>\|t\|) |
| --- | --- | --- | --- | --- | --- |
| (Intercept) | 0.513 | 0.036 | 69.680 | 14.337 | 0.000 |
| Conditionwithin | 0.036 | 0.041 | 48.000 | 0.875 | 0.386 |

Dice within localizer vs. between localizer versions (right-hemisphere wholebrain top 20%), $r^2 = 0.2916$
|  | Estimate | Std. Error | df | t value | Pr(>\|t\|) |
| --- | --- | --- | --- | --- | --- |
| (Intercept) | 0.944 | 0.025 | 61.579 | 38.090 | 0.000 |
| Conditionwithin | -0.008 | 0.026 | 48.000 | -0.312 | 0.756 |

Dice within localizer vs. between localizer versions (right-hemisphere wholebrain top 30%), $r^2 = 0.0911$
|  | Estimate | Std. Error | df | t value | Pr(>\|t\|) |
| --- | --- | --- | --- | --- | --- |
| (Intercept) | 0.966 | 0.003 | 70.960 | 335.496 | 0.000 |
| Conditionwithin | -0.002 | 0.003 | 48.000 | -0.688 | 0.495 |

## SI 3: Information related to Results Section 1.B

### SI 3A: Validation of *hard* > *easy* contrast from the MD localizer

**SI Figure 3A.**
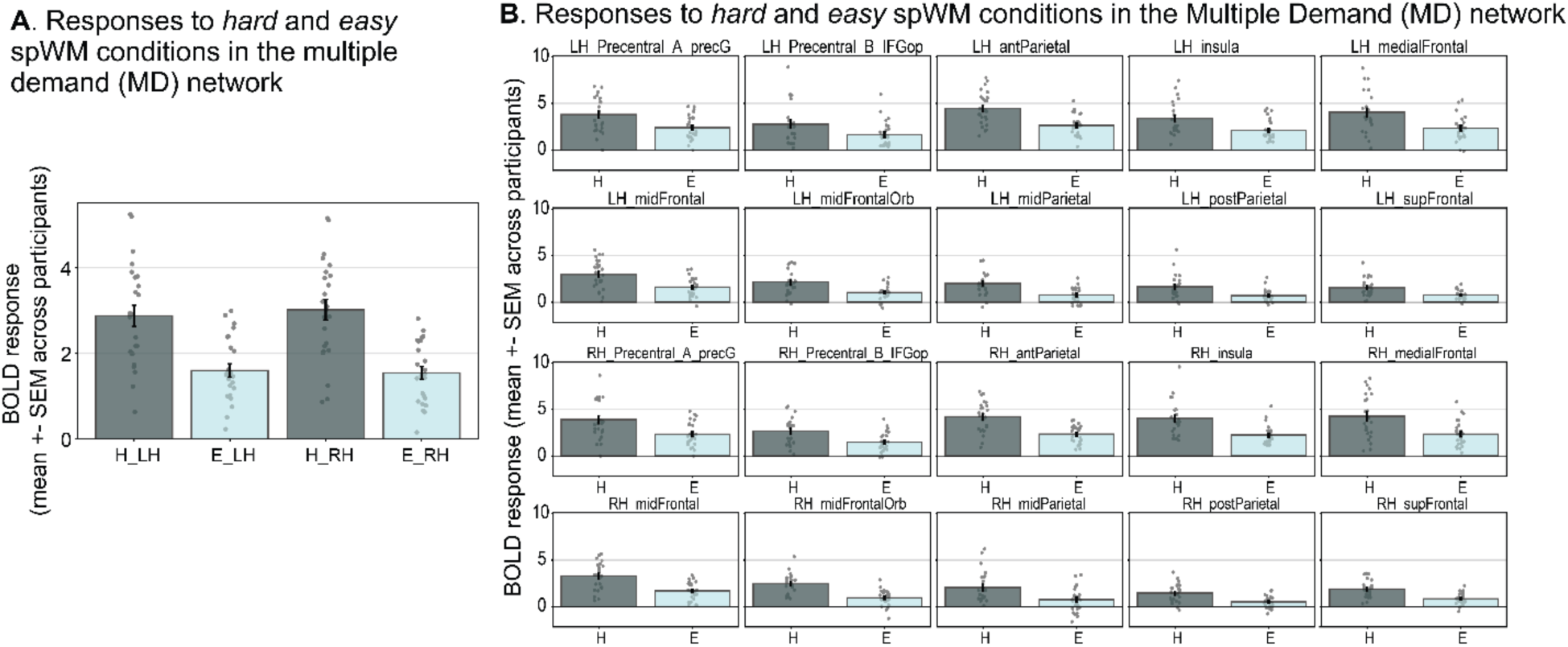
Responses to the hard and easy spatial working memory (spWM) conditions in the MD localizer task. **(A)** Mean response to the MD localizer conditions (H=*hard*, E=*easy*) averaged across ten Multiple Demand (MD) fROIs in each hemisphere. Dots show the mean response across fROIs of each individual participant. Error bars show the standard error of the mean across participants. **(B)** Mean response to the MD localizer conditions for each MD fROI. Dots show the mean response in the particular fROI of each individual participant. Error bars show the standard error of the mean across participants.

### SI 3B: *Sentences* > *nonwords* BOLD response magnitudes are highly correlated across runs for both the standard and speeded language localizer versions

To investigate how stable the *sentences* and *nonwords* BOLD responses were across individual scanning runs, we quantified the average BOLD response magnitudes of the *sentences > nonwords* contrast for each LH language fROI for the odd and even run of each localizer version separately. Note that independent data were used to localize the fROI (i.e., data from the odd run were used to define the fROI, and responses were extracted from the even run, and vice versa).

The correlation between the average *sentence > nonwords* magnitude across LH language fROIs was greater in the standard language localizer than the speeded language localizer (**SI Figure 3B, panel A**). The correlation of the *sentences* > *nonwords* magnitude between odd and even runs across the five language fROIs was r = 0.82 (p<0.0001) for the standard language localizer, and r = 0.57 (p=0.0035) for the speeded language localizer. (Note that without the one outlier participant–bottom right in **SI Figure 3B, panel A**–the correlation was r = 0.83 for the speeded language localizer, p<0.0001).

The odd-even correlation values for individual fROIs were similarly high (**SI Figure 3B, panel B**): The average correlation across the five language fROIs was 0.823 (SD across fROIs: 0.062; five ps<0.001) for the standard language localizer, and 0.704 (SD: 0.083; five ps<0.001) for the speeded language localizer.

**SI Figure 3B.**
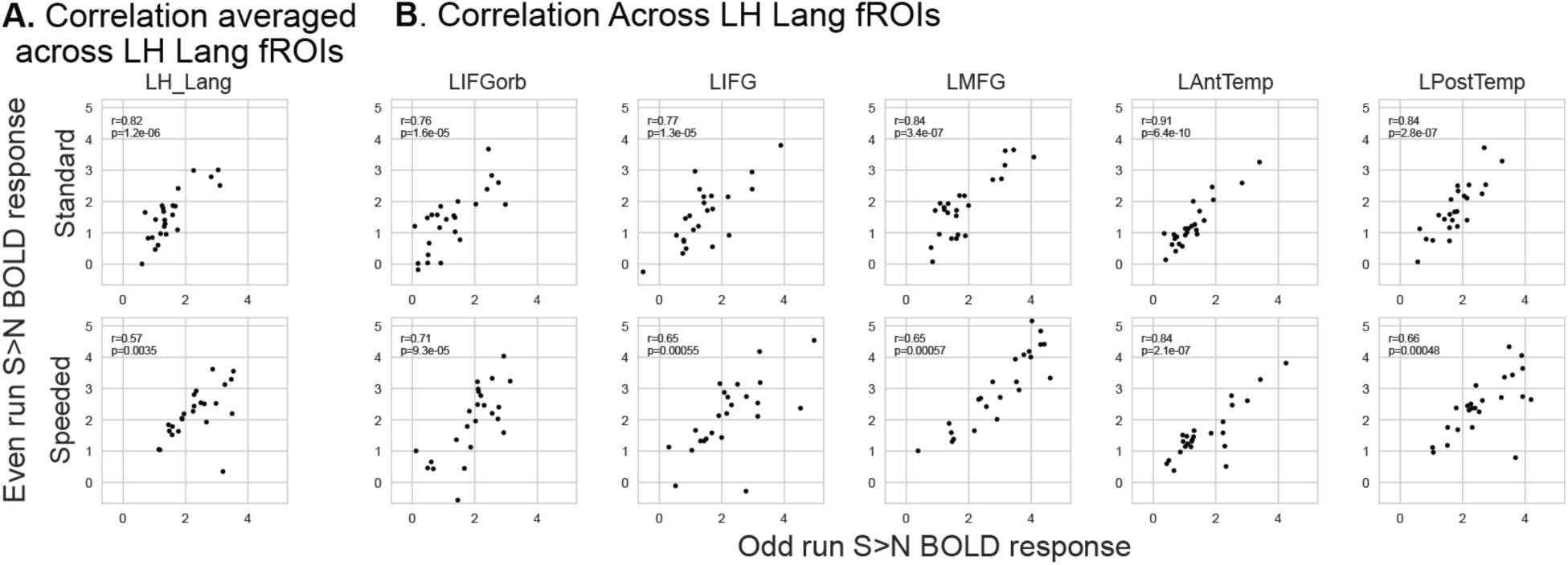
Correlation of *sentences* > *nonwords* BOLD magnitudes within LH language fROIs obtained from odd and even runs. **(A)** Correlation between *sentences > nonwords* BOLD magnitudes (averaged across the five LH language ROIs) of odd (x-axis) and even (y-axis) runs of the standard language localizer (upper row) and speeded language localizer (bottom row). Dots represent the *sentences* > *nonwords* BOLD response for each individual participant (*n*=24). **(B)** Same as in panel A, just for each individual language fROI.

### SI 3C: Consistency of localizers within participants *across* sessions

**SI Figure 3C:**
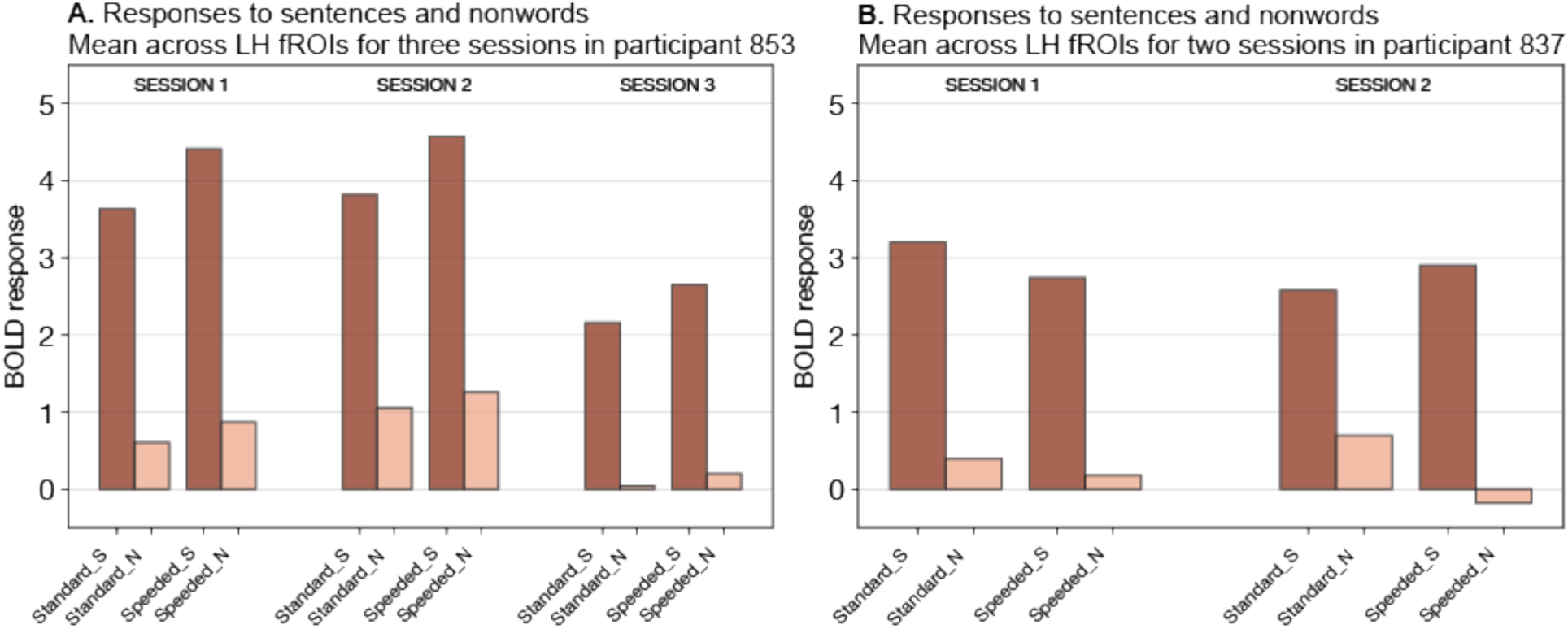
Responses to the *sentences* and *nonwords* conditions in the standard and speeded language localizer for two participants across sessions. Two participants in our dataset completed the two language localizers in different sessions (i.e., different days): One participant completed three sessions (the three sessions were 5 and 7 days apart, **panel A**), another participant completed two sessions (the two sessions were 99 days apart, **panel B**). The mean responses to the language localizer conditions (S=*sentences*, N=*nonwords*) for each of the standard and speeded versions of the localizer task averaged across the five LH language fROIs are shown.

### SI 3D: Statistics tables for Results Section 1.B (responses to language)

In the tables below, “BOLD response” denotes the BOLD response magnitude for the given condition (*sentences, nonwords,* or *sentences > nonwords;* note that “language” denotes both *sentences* and *nonwords* responses). “condition” denotes the *sentences* and *nonwords* conditions in the LMEs where they are modeled together. “version” denotes the language localizer version, either standard or speeded. “participant” denotes each of the *n*=24 participants. “fROI” denotes each of the five LH fROIs.

|  | Estimate | Std. Error | df | t value | Pr(>\|t\|) |
| --- | --- | --- | --- | --- | --- |
| (Intercept) | 0.442 | 0.270 | 7.587 | 1.636 | 0.142 |
| ConditionS | 1.897 | 0.080 | 451.954 | 23.570 | 0.000 |
| VersionSpeeded | -0.068 | 0.080 | 451.954 | -0.851 | 0.395 |

(ii) *BOLD response language* ~ *condition* + *version* + *condition:version* + (*I*|*participant*) + (*I*|*fROI*) Standard vs Speeded With Interaction (LHLang), $r^2 = 0.6671$
|  | Estimate | Std. Error | df | t value | Pr(>\|t\|) |
| --- | --- | --- | --- | --- | --- |
| (Intercept) | 0.623 | 0.273 | 7.895 | 2.282 | 0.052 |
| ConditionS | 1.535 | 0.111 | 451.955 | 13.801 | 0.000 |
| VersionSpeeded | -0.430 | 0.111 | 451.955 | -3.866 | 0.000 |
| ConditionS:VersionSpeeded | 0.723 | 0.157 | 451.955 | 4.597 | 0.000 |

(iii) *BOLD response sentences > nonwords* ~ *version* + (*I*|*participant*) + (*I*|*fROI*) Standard vs Speeded S-N (LHLang), $r^2 = 0.5214$
|  | Estimate | Std. Error | df | t value | Pr(>\|t\|) |
| --- | --- | --- | --- | --- | --- |
| (Intercept) | 1.535 | 0.204 | 10.980 | 7.521 | 0.000 |
| VersionSpeeded | 0.723 | 0.093 | 211.888 | 7.817 | 0.000 |

(iv) *BOLD response sentences* ~ *version* + (*I*|*participant*) + (*I*|*fROI*) Standard vs Speeded S (LHLang), $r^2 = 0.546$
|  | Estimate | Std. Error | df | t value | Pr(>\|t\|) |
| --- | --- | --- | --- | --- | --- |
| (Intercept) | 2.158 | 0.361 | 7.700 | 5.978 | 0.000 |
| VersionSpeeded | 0.293 | 0.122 | 211.940 | 2.407 | 0.017 |

(v) *BOLD response nonwords* ~ *version* + (*I*|*participant*) + (*I*|*fROI*) Standard vs Speeded N (LHLang), $r^2 = 0.3951$
|  | Estimate | Std. Error | df | t value | Pr(>\|t\|) |
| --- | --- | --- | --- | --- | --- |
| (Intercept) | 0.623 | 0.182 | 7.397 | 3.418 | 0.010 |
| VersionSpeeded | -0.430 | 0.084 | 211.884 | -5.106 | 0.000 |

### SI 3E: Language BOLD responses for right hemisphere fROIs

**SI Figure 3E:**
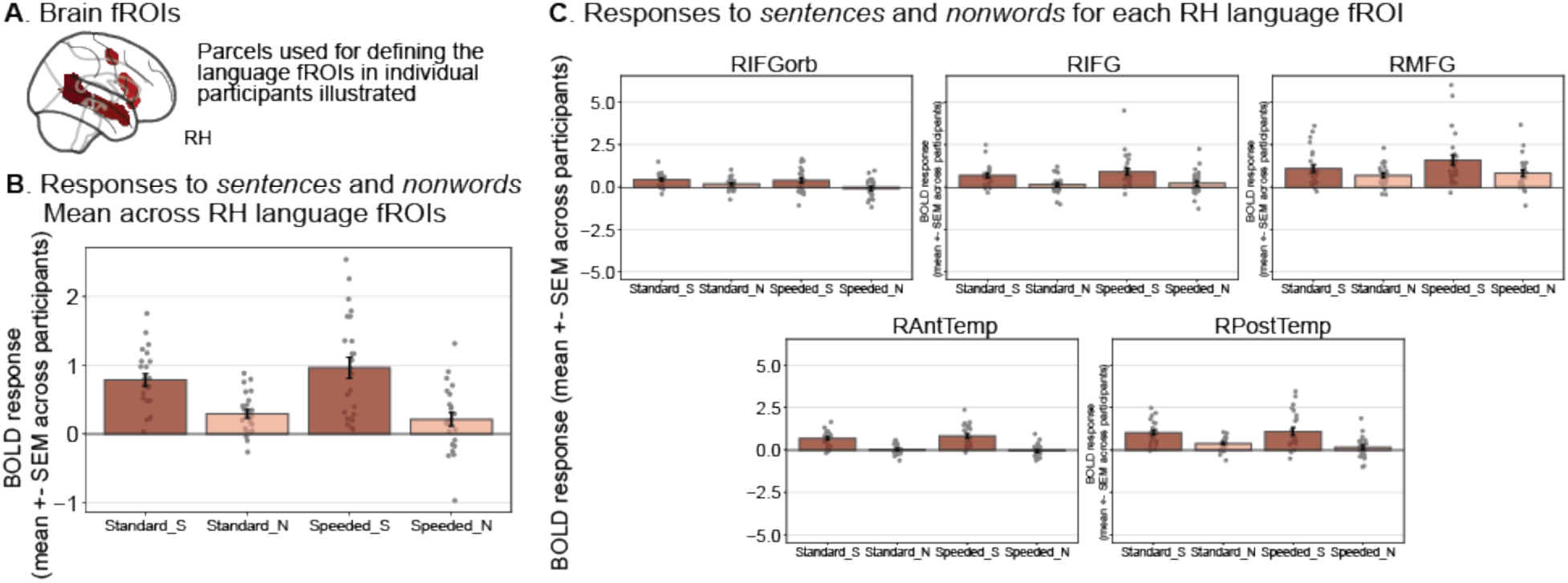
Responses to the language localizer conditions (*sentences* and *nonwords*) for the standard and speeded language localizers in right hemisphere (RH) language fROIs. **(A)** We defined the RH language fROIs as the most language-responsive voxels (top 10%) within the borders of the five anatomical parcels (see <u>Methods; Extraction of fMRI BOLD responses</u>) for each participant, and measured the BOLD response magnitude in these fROIs in a cross-validated manner (see <u>Methods; Definition of fROIs</u>). **(B)** Mean BOLD response to the language localizer conditions (S=*sentences*, N=*nonwords*) for both the standard and speeded localizer versions averaged across the five RH language fROIs. **(C)** Mean BOLD response to the localizer conditions for each RH language fROI. In both panels, dots show the mean response of each individual participant. Error bars show the standard error of the mean across participants.

The statistics tables accompanying SI Figure 3E are found below.

|  | Estimate | Std. Error | df | t value | Pr(>\|t\|) |
| --- | --- | --- | --- | --- | --- |
| (Intercept) | 0.228 | 0.146 | 8.359 | 1.555 | 0.157 |
| ConditionS | 0.623 | 0.065 | 451.888 | 9.617 | 0.000 |
| VersionSpeeded | 0.047 | 0.065 | 451.888 | 0.732 | 0.465 |

(ii) *BOLD response RH language ~ condition + version + condition:version + (I|participant) + (I|fROI)* Standard vs Speeded With Interaction (RHLang), $r^2 = 0.3332$
|  | Estimate | Std. Error | df | t value | Pr(>\|t\|) |
| --- | --- | --- | --- | --- | --- |
| (Intercept) | 0.291 | 0.150 | 9.180 | 1.942 | 0.083 |
| ConditionS | 0.496 | 0.091 | 451.889 | 5.438 | 0.000 |
| VersionSpeeded | -0.079 | 0.091 | 451.889 | -0.872 | 0.384 |
| ConditionS:VersionSpeeded | 0.254 | 0.129 | 451.889 | 1.968 | 0.050 |

(iii) *BOLD response RH sentences>nonwords delta ~ version + (I|participant) + (I|fROI)* Standard vs Speeded S-N (RHLang), $r^2 = 0.3587$
|  | Estimate | Std. Error | df | t value | Pr(>\|t\|) |
| --- | --- | --- | --- | --- | --- |
| (Intercept) | 0.496 | 0.105 | 17.126 | 4.717 | 0.000 |
| VersionSpeeded | 0.254 | 0.067 | 211.758 | 3.776 | 0.000 |

(iv) *BOLD response RH sentences ~ version + (I|participant) + (I|fROI)* Standard vs Speeded S (RHLang), $r^2 = 0.2493$
|  | Estimate | Std. Error | df | t value | Pr(>\|t\|) |
| --- | --- | --- | --- | --- | --- |
| (Intercept) | 0.787 | 0.168 | 9.317 | 4.678 | 0.001 |
| VersionSpeeded | 0.174 | 0.106 | 211.768 | 1.651 | 0.100 |

(v) *BOLD response RH nonwords ~ version + (I|participant) + (I|fROI)* Standard vs Speeded N (RHLang), $r^2 = 0.2648$
|  | Estimate | Std. Error | df | t value | Pr(>\|t\|) |
| --- | --- | --- | --- | --- | --- |
| (Intercept) | 0.291 | 0.134 | 7.002 | 2.175 | 0.066 |
| VersionSpeeded | -0.079 | 0.071 | 211.842 | -1.118 | 0.265 |

### SI 3F: Statistics tables for Results Section 1.B (responses to working memory task)

In the tables below, “BOLD response” denotes the BOLD response magnitude for the given condition (*hard, easy*). “version” denotes the language localizer version, either standard or speeded. “participant” denotes each of the *n*=24 participants. “fROI” denotes each of the five LH fROIs.

|  | Estimate | Std. Error | df | t value | Pr(>\|t\|) |
| --- | --- | --- | --- | --- | --- |
| (Intercept) | -0.383 | 0.284 | 9.648 | -1.347 | 0.209 |
| VersionSpeeded | 0.101 | 0.104 | 211.929 | 0.973 | 0.332 |

ii) *BOLD response easy* ∼ version + ( $1|$ participant) + ( $1|$ fROI) Standard vs Speeded E (LHLang), $r^2 = 0.4984$
|  | Estimate | Std. Error | df | t value | Pr(>\|t\|) |
| --- | --- | --- | --- | --- | --- |
| (Intercept) | -0.054 | 0.156 | 13.538 | -0.343 | 0.737 |
| VersionSpeeded | 0.096 | 0.070 | 211.891 | 1.361 | 0.175 |

### SI 3G: Language responses in extended language network fROIs

**SI Figure 3G.**
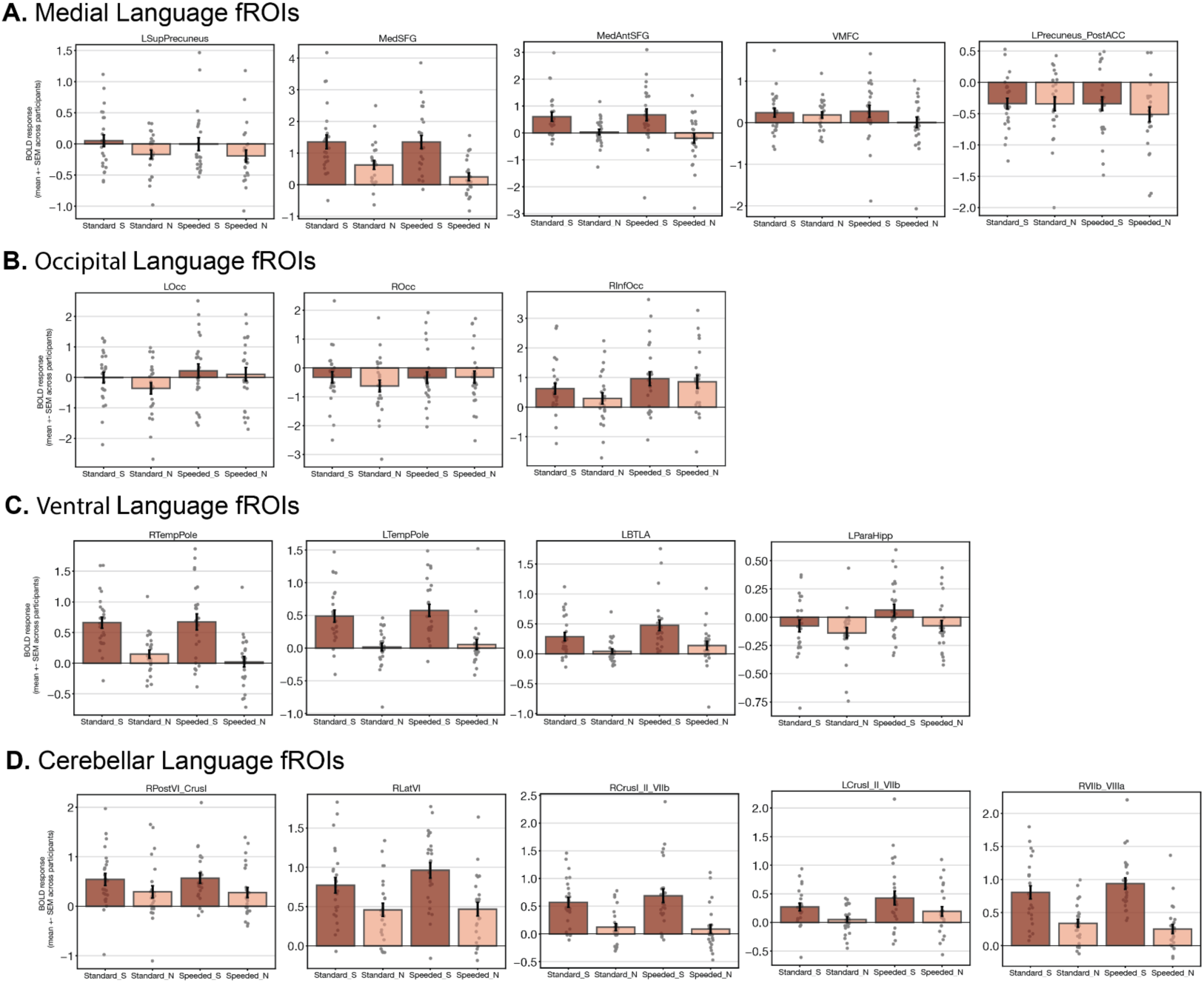
Responses to the standard and speeded localizer tasks in the extended language network. Mean BOLD response to the language localizer conditions (S=sentences, N=nonwords) for each of the standard and speeded versions of the localizer task for each fROI in the **(A)** Medial, **(B)** Occipital, **(C)** Ventral, and **(D)** Cerebellar regions of the extended language network as described by Wolna et al. (2025). Dots show the mean response across fROIs of each individual participant. Error bars show the standard error of the mean across participants.

### SI 3H: Language responses in DMN fROIs

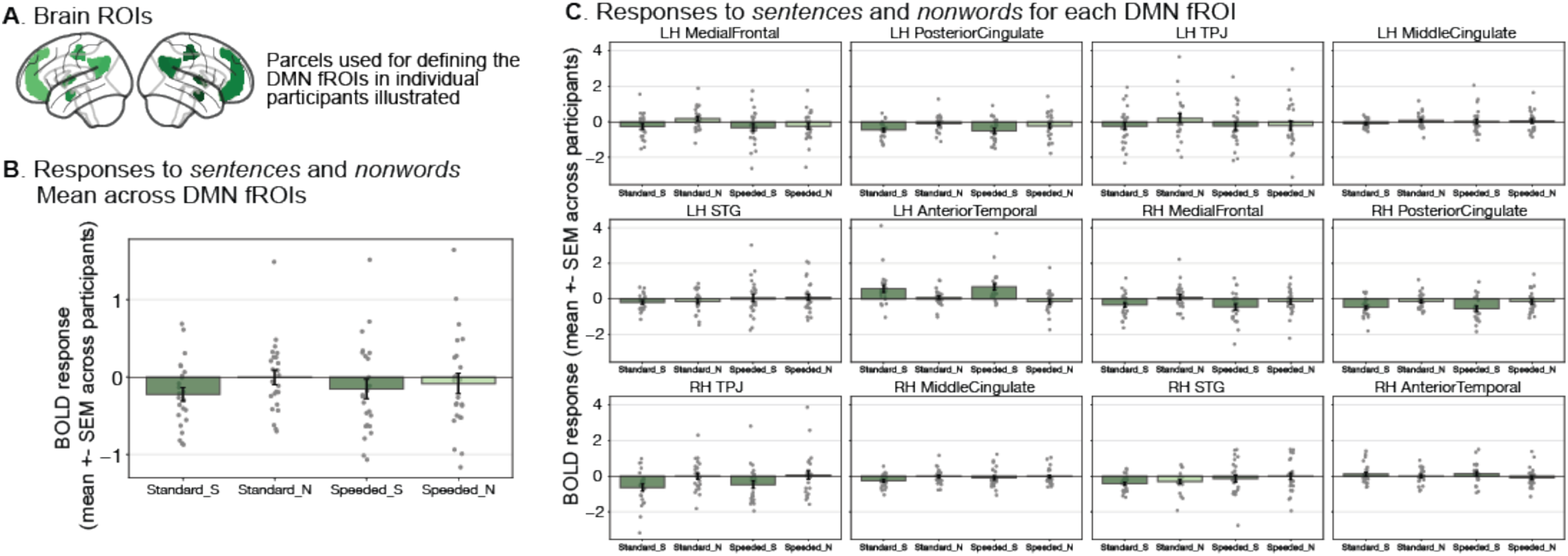

**(A)** To define the DMN fROIs, we used a set of 12 parcels (6 in each hemisphere) derived from a group-level probabilistic activation overlap map for the *easy > hard* spatial working memory contrast (Fedorenko et al., 2013) in 197 independent participants. The *easy > hard* contrast has been shown to robustly activate DMN regions, in line with prior work using similar tasks and contrasts (Leech et al., 2011; Mineroff, Blank et al., 2018). The parcels included symmetrical regions in the posterior cingulate cortex, four medial frontal regions, the precuneus, and temporoparietal junction. We used the *easy > hard* spatial working memory (WM) contrast to identify the top 10% most responsive voxels within these parcels, and extracted responses to sentences and nonwords conditions within these fROIs for both the standard and speeded language localizer versions. We measured the BOLD response magnitude to the language localizer conditions in these fROIs in a cross-validated manner (see <u>Methods; Definition of fROIs</u>).

**(B)** Mean BOLD response to the language localizer conditions (S=sentences, N=nonwords) for both the standard and speeded localizer versions averaged across the DMN fROIs. On average, the DMN parcels demonstrated a *nonwords > sentences* effect (*sentences > nonwords*, ꞵ=-0.218, t=-3.913, p<0.0001).

**(C)** Mean BOLD response to the localizer conditions for each DMN fROIs. In both panels, dots show the mean response of each individual participant. Error bars show the standard error of the mean across participants. With the exception of the left anterior-temporal region, all DMN regions demonstrated a *nonwords > sentences* response. Notably, this left anterior-temporal region lies within the left anterior-temporal region of the language parcel, and hence the positive *sentences > nonwords* response is likely explained by language-selective voxels in this region.

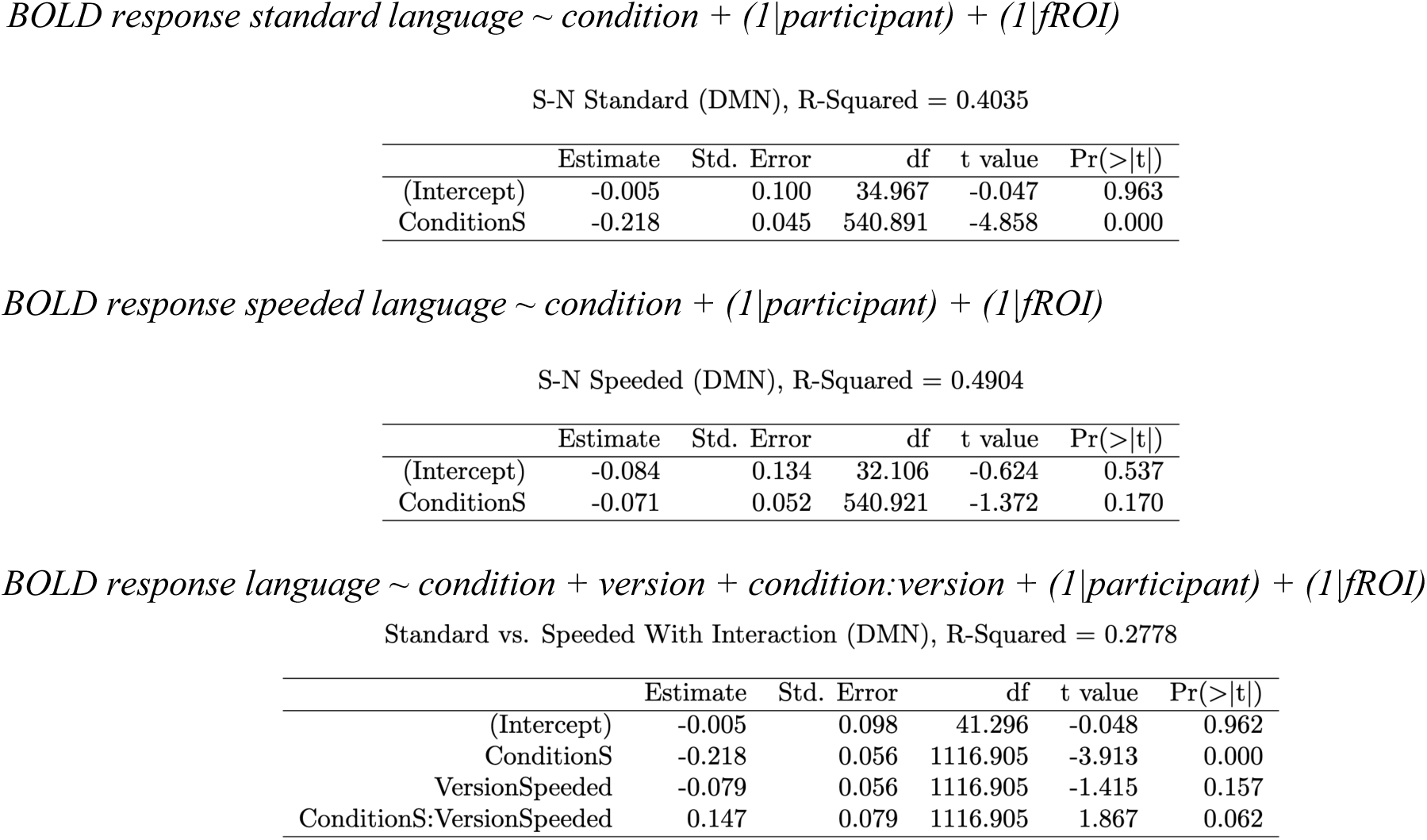

|  | Estimate | Std. Error | df | t value | Pr(>\|t\|) |
| --- | --- | --- | --- | --- | --- |
| (Intercept) | -0.005 | 0.100 | 34.967 | -0.047 | 0.963 |
| ConditionS | -0.218 | 0.045 | 540.891 | -4.858 | 0.000 |

*BOLD response speeded language ~ condition + (1|participant) + (1|fROI)* S-N Speeded (DMN), R-Squared = 0.4904
|  | Estimate | Std. Error | df | t value | Pr(>\|t\|) |
| --- | --- | --- | --- | --- | --- |
| (Intercept) | -0.084 | 0.134 | 32.106 | -0.624 | 0.537 |
| ConditionS | -0.071 | 0.052 | 540.921 | -1.372 | 0.170 |

*BOLD response language ~ condition + version + condition:version + (1|participant) + (1|fROI)* Standard vs. Speeded With Interaction (DMN), R-Squared = 0.2778
|  | Estimate | Std. Error | df | t value | Pr(>\|t\|) |
| --- | --- | --- | --- | --- | --- |
| (Intercept) | -0.005 | 0.098 | 41.296 | -0.048 | 0.962 |
| ConditionS | -0.218 | 0.056 | 1116.905 | -3.913 | 0.000 |
| VersionSpeeded | -0.079 | 0.056 | 1116.905 | -1.415 | 0.157 |
| ConditionS:VersionSpeeded | 0.147 | 0.079 | 1116.905 | 1.867 | 0.062 |

## SI 4: Information related to Results Section 2 (Multiple Demand network)

### SI 4A: MD responses to the language localizer versions across fROIs

**SI Figure 4A.**
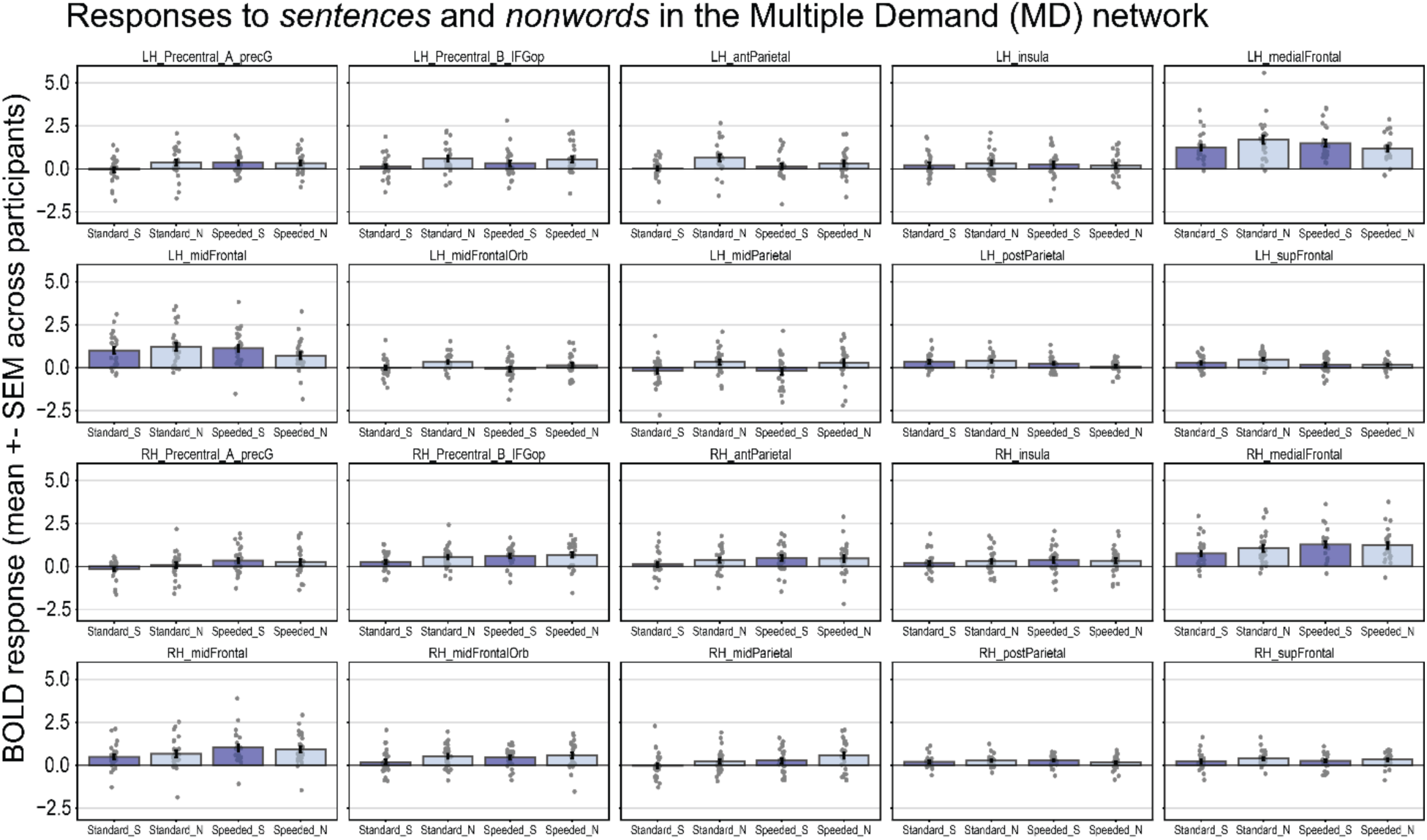
Responses to the standard and speeded localizer tasks in the Multiple Demand Region across all 10 left and right hemispheric MD fROIs. Mean BOLD response to the language localizer conditions (S=*sentences,* N=*nonwords*) for each of the standard and speeded versions of the localizer task for each Multiple Demand (MD) fROI in the left and right hemispheres. Dots show the mean response across fROIs of each individual participant. Error bars show the standard error of the mean across participants.

### SI 4B: Statistics tables for Results Section 2

In the tables below, “BOLD response” denotes the BOLD response magnitude for the given condition (*sentences, nonwords*; note that “language” denotes both *sentences* and *nonwords* responses). “condition” denotes the *sentences* and *nonwords* conditions in the LMEs where they are modeled together. “version” denotes the language localizer version, either standard or speeded. “participant” denotes each of the *n*=24 participants. “fROI” denotes each of the twenty LH/RH MD fROIs.

|  | Estimate | Std. Error | df | t value | Pr(>\|t\|) |
| --- | --- | --- | --- | --- | --- |
| (Intercept) | 0.541 | 0.125 | 42.783 | 4.325 | 0.000 |
| ConditionS | -0.275 | 0.036 | 916.956 | -7.547 | 0.000 |

*BOLD response speeded language* ∼ condition + (1|participant) + (1|fROI) S-N Speeded (MD), $r^2 = 0.44$
|  | Estimate | Std. Error | df | t value | Pr(>\|t\|) |
| --- | --- | --- | --- | --- | --- |
| (Intercept) | 0.477 | 0.123 | 42.998 | 3.864 | 0.000 |
| ConditionS | -0.012 | 0.041 | 916.942 | -0.283 | 0.777 |

*BOLD response language* ∼ condition + version + condition:version + (1|participant) + (1|fROI) Standard vs. Speeded With Interaction (MD), $r^2 = 0.3504$
|  | Estimate | Std. Error | df | t value | Pr(>\|t\|) |
| --- | --- | --- | --- | --- | --- |
| (Intercept) | 0.541 | 0.108 | 45.171 | 5.002 | 0.000 |
| ConditionS | -0.275 | 0.044 | 1876.956 | -6.328 | 0.000 |
| VersionSpeeded | -0.064 | 0.044 | 1876.956 | -1.470 | 0.142 |
| ConditionS:VersionSpeeded | 0.264 | 0.062 | 1876.956 | 4.285 | 0.000 |

*BOLD response sentences* ∼ version + (1|participant) + (1|fROI) Standard vs. Speeded S (MD), $r^2 = 0.3263$
|  | Estimate | Std. Error | df | t value | Pr(>\|t\|) |
| --- | --- | --- | --- | --- | --- |
| (Intercept) | 0.266 | 0.104 | 36.377 | 2.549 | 0.015 |
| VersionSpeeded | 0.200 | 0.044 | 916.907 | 4.584 | 0.000 |

*BOLD response nonwords* ∼ version + (1|participant) + (1|fROI) Standard vs. Speeded N (MD), $r^2 = 0.4024$
|  | Estimate | Std. Error | df | t value | Pr(>\|t\|) |
| --- | --- | --- | --- | --- | --- |
| (Intercept) | 0.541 | 0.117 | 42.660 | 4.634 | 0.000 |
| VersionSpeeded | -0.064 | 0.042 | 916.932 | -1.519 | 0.129 |

## References

Adank, P., Davis, M. H., & Hagoort, P. (2012). Neural dissociation in processing noise and accent in spoken language comprehension. Neuropsychologia, 50(1), 77–84. 10.1016/j.neuropsychologia.2011.10.024

Adank, P., & Janse, E. (2010). Comprehension of a novel accent by young and older listeners. Psychology and Aging, 25(3), 736–740. 10.1037/a0020054

Allen, E. J., St-Yves, G., Wu, Y., Breedlove, J. L., Prince, J. S., Dowdle, L. T., Nau, M., Caron, B., Pestilli, F., Charest, I., Hutchinson, J. B., Naselaris, T., & Kay, K. (2022). A massive 7T fMRI dataset to bridge cognitive neuroscience and artificial intelligence. Nature Neuroscience, 25(1), Article 1. 10.1038/s41593-021-00962-x

Amalric, M., & Dehaene, S. (2019). A distinct cortical network for mathematical knowledge in the human brain. NeuroImage, 189, 19–31. 10.1016/j.neuroimage.2019.01.001

Ashburner, J., & Friston, K. J. (2005). Unified segmentation. NeuroImage, 26(3), 839–851. 10.1016/j.neuroimage.2005.02.018

Assem, M., Blank, I. A., Mineroff, Z., Ademoğlu, A., & Fedorenko, E. (2020a). Activity in the fronto-parietal multiple-demand network is robustly associated with individual differences in working memory and fluid intelligence. Cortex, 131, 1–16. 10.1016/j.cortex.2020.06.013

Assem, M., Glasser, M. F., Van Essen, D. C., & Duncan, J. (2020b). A Domain-General Cognitive Core Defined in Multimodally Parcellated Human Cortex. Cerebral Cortex, 30(8), 4361–4380. 10.1093/cercor/bhaa023

Baker, C. I., Liu, J., Wald, L. L., Kwong, K. K., Benner, T., & Kanwisher, N. (2007). Visual word processing and experiential origins of functional selectivity in human extrastriate cortex. Proceedings of the National Academy of Sciences, 104(21), 9087–9092. 10.1073/pnas.0703300104

Banks, B., Gowen, E., Munro, K. J., & Adank, P. (2015). Cognitive predictors of perceptual adaptation to accented speech. The Journal of the Acoustical Society of America, 137(4), 2015–2024. 10.1121/1.4916265

Bashivan, P., Kar, K., & DiCarlo, J. J. (2019). Neural population control via deep image synthesis. Science, 364(6439), eaav9436. 10.1126/science.aav9436

Bates, D., Mächler, M., Bolker, B., & Walker, S. (2015). Fitting Linear Mixed-Effects Models Using lme4. Journal of Statistical Software, 67, 1–48. 10.18637/jss.v067.i01

Bates, E., Wilson, S. M., Saygin, A. P., Dick, F., Sereno, M. I., Knight, R. T., & Dronkers, N. F. (2003). Voxel-based lesion-symptom mapping. Nature Neuroscience, 6(5), 448–450. 10.1038/nn1050

Bedny, M., Pascual-Leone, A., Dodell-Feder, D., Fedorenko, E., & Saxe, R. (2011). Language processing in the occipital cortex of congenitally blind adults. Proceedings of the National Academy of Sciences, 108(11), 4429–4434. 10.1073/pnas.1014818108

Belin, P., Zatorre, R. J., & Ahad, P. (2002). Human temporal-lobe response to vocal sounds. Cognitive Brain Research, 13(1), 17–26. 10.1016/S0926-6410(01)00084-2

Benjamin, C. F., & Gaab, N. (2012). What’s the story? The tale of reading fluency told at speed. Human brain mapping, 33(11), 2572–2585.

Billot, A. (2023). Neuroplasticity mechanisms in post-stroke aphasia: Investigating the differential role of the domain-general multiple demand and language networks [Doctoral dissertation]. Boston University.

Billot, A., Jhingan, N., Varkanitsa, M., Blank, I., Ryskin, R., Kiran, S., & Fedorenko, E. (2024). The language network ages well: Preserved selectivity, lateralization, and within-network functional synchronization in older brains. bioRxiv, doi: 10.1101/2024.10.23.619954.

Binder, J. R., Frost, J. A., Hammeke, T. A., Cox, R. W., Rao, S. M., & Prieto, T. (1997). Human brain language areas identified by functional magnetic resonance imaging. The Journal of Neuroscience: The Official Journal of the Society for Neuroscience, 17(1), 353–362.

Bird, S., & Loper, E. (2004). NLTK: The Natural Language Toolkit. Proceedings of the ACL Interactive Poster and Demonstration Sessions, 214–217. https://aclanthology.org/P04-3031

Blank, I. A., & Fedorenko, E. (2017). Domain-General Brain Regions Do Not Track Linguistic Input as Closely as Language-Selective Regions. The Journal of Neuroscience, 37(41), 9999–10011. 10.1523/JNEUROSCI.3642-16.2017

Blank, I., Balewski, Z., Mahowald, K., & Fedorenko, E. (2016). Syntactic processing is distributed across the language system. NeuroImage, 127, 307–323. 10.1016/j.neuroimage.2015.11.069

Blank, I., Kanwisher, N., & Fedorenko, E. (2014). A functional dissociation between language and multiple-demand systems revealed in patterns of BOLD signal fluctuations. Journal of Neurophysiology, 112(5), 1105–1118. 10.1152/jn.00884.2013

Braga, R. M., DiNicola, L. M., Becker, H. C., & Buckner, R. L. (2020). Situating the left-lateralized language network in the broader organization of multiple specialized large-scale distributed networks. Journal of Neurophysiology, 124(5), 1415–1448. 10.1152/jn.00753.2019

Brett, M., Johnsrude, I. S., & Owen, A. M. (2002). The problem of functional localization in the human brain. Nature Reviews. Neuroscience, 3(3), 243–249. 10.1038/nrn756

Bugatus, L., Weiner, K. S., & Grill-Spector, K. (2017). Task alters category representations in prefrontal but not high-level visual cortex. Neuroimage, 155, 437–449.

Chen, E. M., Kamps, F. S., & Saxe, R. R. (2024). An open dataset of functional MRI responses to egocentric navigation through natural scenes. Cognitive Computational Neuroscience.

Chen, X., Affourtit, J., Ryskin, R., Regev, T. I., Norman-Haignere, S., Jouravlev, O., Malik-Moraleda, S., Kean, H., Varley, R., & Fedorenko, E. (2023). The human language system, including its inferior frontal component in “Broca’s area,” does not support music perception. Cerebral Cortex, bhad087. 10.1093/cercor/bhad087

Christodoulou, J. A., Del Tufo, S. N., Lymberis, J., Saxler, P. K., Ghosh, S. S., Triantafyllou, C., Whitfield-Gabrieli, S., & Gabrieli, J. D. (2014). Brain bases of reading fluency in typical reading and impaired fluency in dyslexia. PloS one, 9(7), e100552

Clercq, P. D., Gonsalves, A. R., Gerrits, R., & Vandermosten, M. (2024). Individualized functional localization of the language and multiple demand network in chronic post-stroke aphasia (p. 2024.01.12.575350). bioRxiv. 10.1101/2024.01.12.575350

Deen, B., Koldewyn, K., Kanwisher, N., & Saxe, R. (2015). Functional Organization of Social Perception and Cognition in the Superior Temporal Sulcus. Cerebral Cortex (New York, N.Y.: 1991), 25(11), 4596–4609. 10.1093/cercor/bhv111

Diachek, E., Blank, I., Siegelman, M., Affourtit, J., & Fedorenko, E. (2020). The domain-general multiple demand (MD) network does not support core aspects of language comprehension: A large-scale fMRI investigation. Journal of Neuroscience. 10.1523/JNEUROSCI.2036-19.2020

Dice, L. R. (1945). Measures of the Amount of Ecologic Association Between Species. Ecology, 26(3), 297–302. 10.2307/1932409

DiNicola, L. M., Ariyo, O. I., & Buckner, R. L. (2023). Functional specialization of parallel distributed networks revealed by analysis of trial-to-trial variation in processing demands. Journal of neurophysiology, 129(1), 17–40.

Dodell-Feder, D., Koster-Hale, J., Bedny, M., & Saxe, R. (2011). fMRI item analysis in a theory of mind task. NeuroImage, 55(2), 705–712. 10.1016/j.neuroimage.2010.12.040

Downing, P. E., Jiang, Y., Shuman, M., & Kanwisher, N. (2001). A cortical area selective for visual processing of the human body. Science (New York, N.Y.), 293(5539), 2470–2473. 10.1126/science.1063414

Du, J., DiNicola, L. M., Angeli, P. A., Saadon-Grosman, N., Sun, W., Kaiser, S., Ladopoulou, J., Xue, A., Yeo, B. T. T., Eldaief, M. C., & Buckner, R. L. (2024). Organization of the human cerebral cortex estimated within individuals: Networks, global topography, and function. Journal of Neurophysiology, 131(6), 1014–1082. 10.1152/jn.00308.2023

Du, J., Tripathi, V., Elliott, M. L., Ladopoulou, J., Sun, W., Eldaief, M. C., & Buckner, R. L. (2025). Within-individual precision mapping of brain networks exclusively using task data. Neuron, 113(23), 4069–4083.

Duncan, J. (2010). The multiple-demand (MD) system of the primate brain: Mental programs for intelligent behaviour. Trends in Cognitive Sciences, 14(4), 172–179. 10.1016/j.tics.2010.01.004

Duncan, J., Assem, M., & Shashidhara, S. (2020). Integrated Intelligence from Distributed Brain Activity. Trends in Cognitive Sciences, 24(10), 838–852. 10.1016/j.tics.2020.06.012

Duncan, J., & Owen, A. M. (2000). Common regions of the human frontal lobe recruited by diverse cognitive demands. Trends in Neurosciences, 23(10), 475–483. 10.1016/s0166-2236(00)01633-7

Duncan, J., Schramm, M., Thompson, R., & Dumontheil, I. (2012). Task rules, working memory, and fluid intelligence. Psychonomic Bulletin & Review, 19(5), 864–870. 10.3758/s13423-012-0225-y

Epstein, R., & Kanwisher, N. (1998). A cortical representation of the local visual environment. Nature, 392(6676), 598–601. 10.1038/33402

Fedorenko, E., Behr, M. K., & Kanwisher, N. (2011). Functional specificity for high-level linguistic processing in the human brain. Proceedings of the National Academy of Sciences, 108(39), 16428–16433. 10.1073/pnas.1112937108

Fedorenko, E., & Blank, I. A. (2020). Broca’s Area Is Not a Natural Kind. Trends in Cognitive Sciences, 24(4), 270–284. 10.1016/j.tics.2020.01.001

Fedorenko, E., Blank, I. A., Siegelman, M., & Mineroff, Z. (2020). Lack of selectivity for syntax relative to word meanings throughout the language network. Cognition, 203, 104348. 10.1016/j.cognition.2020.104348

Fedorenko, E., Duncan, J., & Kanwisher, N. (2012). Language-Selective and Domain-General Regions Lie Side by Side within Broca’s Area. Current Biology, 22(21), 2059–2062. 10.1016/j.cub.2012.09.011

Fedorenko, E., Duncan, J., & Kanwisher, N. (2013). Broad domain generality in focal regions of frontal and parietal cortex. Proceedings of the National Academy of Sciences, 110(41), 16616–16621. 10.1073/pnas.1315235110

Fedorenko, E., Hsieh, P.-J., Nieto-Castañón, A., Whitfield-Gabrieli, S., & Kanwisher, N. (2010). New method for fMRI investigations of language: Defining ROIs functionally in individual subjects. Journal of Neurophysiology, 104(2), 1177–1194. 10.1152/jn.00032.2010

Fedorenko, E., Ivanova, A. A., & Regev, T. I. (2024). The language network as a natural kind within the broader landscape of the human brain. Nature Reviews Neuroscience, 25(5), 289–312. 10.1038/s41583-024-00802-4

Fedorenko, E., & Shain, C. (2021). Similarity of computations across domains does not imply shared implementation: The case of language comprehension. Current Directions in Psychological Science, 30(6), 526–534. 10.1177/09637214211046955

Fernandino, L., & Binder, J. R. (2024). How does the “default mode” network contribute to semantic cognition?. Brain and language, 252, 105405.

Fischer, J., Mikhael, J. G., Tenenbaum, J. B., & Kanwisher, N. (2016). Functional neuroanatomy of intuitive physical inference. Proceedings of the National Academy of Sciences, 113(34), E5072–E5081. 10.1073/pnas.1610344113

Fischl, B., Rajendran, N., Busa, E., Augustinack, J., Hinds, O., Yeo, B. T. T., Mohlberg, H., Amunts, K., & Zilles, K. (2008). Cortical folding patterns and predicting cytoarchitecture. Cerebral Cortex (New York, N.Y.: 1991), 18(8), 1973–1980. 10.1093/cercor/bhm225

Forster, K. I. (1970). Visual perception of rapidly presented word sequences of varying complexity. Perception & Psychophysics, 8(4), 215–221. 10.3758/BF03210208

Francis, W. N., & Kucera, H. (1964). A Standard Corpus of Present-Day Edited American English, for use with Digital Computers (Brown) [dataset]. https://varieng.helsinki.fi/CoRD/corpora/BROWN/

Fridriksson, J., den Ouden, D.-B., Hillis, A. E., Hickok, G., Rorden, C., Basilakos, A., Yourganov, G., & Bonilha, L. (2018). Anatomy of aphasia revisited. Brain: A Journal of Neurology, 141(3), 848–862. 10.1093/brain/awx363

Friederici, A. D. (2012). The cortical language circuit: From auditory perception to sentence comprehension. Trends in Cognitive Sciences, 16(5), 262–268. 10.1016/j.tics.2012.04.001

Friston, Karl. J., Ashburner, J., Frith, C. D., Poline, J.-B., Heather, J. D., & Frackowiak, R. S. J. (1995). Spatial registration and normalization of images. Human Brain Mapping, 3(3), 165–189. 10.1002/hbm.460030303

Frost, M. A., & Goebel, R. (2012). Measuring structural-functional correspondence: Spatial variability of specialised brain regions after macro-anatomical alignment. NeuroImage, 59(2), 1369–1381. 10.1016/j.neuroimage.2011.08.035

Gao, R., Cheung, C., Siegelman, M., Pongos, A. L. A., Kean, H. H., Tanner, A., Fedorenko, E., & Ivanova, A. A. (2025). The language network responds robustly to sentences across diverse tasks (p. 2025.12.02.691902). bioRxiv.

Goodglass, H. (1993). Understanding Aphasia. ACADEMIC PressINC.

Gordon, E. M., Laumann, T. O., Gilmore, A. W., Newbold, D. J., Greene, D. J., Berg, J. J., Ortega, M., Hoyt-Drazen, C., Gratton, C., Sun, H., Hampton, J. M., Coalson, R. S., Nguyen, A. L., McDermott, K. B., Shimony, J. S., Snyder, A. Z., Schlaggar, B. L., Petersen, S. E., Nelson, S. M., & Dosenbach, N. U. F. (2017). Precision Functional Mapping of Individual Human Brains. Neuron, 95(4), 791–807.e7. 10.1016/j.neuron.2017.07.011

Gratton, C., & Braga, R. M. (2021). Editorial overview: Deep imaging of the individual brain: past, practice, and promise. Current Opinion in Behavioral Sciences, 40, iii–vi. 10.1016/j.cobeha.2021.06.011

Grill-Spector, K., Henson, R., & Martin, A. (2006). Repetition and the brain: neural models of stimulus-specific effects. Trends in cognitive sciences, 10(1), 14–23

Gu, Z., Jamison, K., Sabuncu, M. R., & Kuceyeski, A. (2023). Modulating human brain responses via optimal natural image selection and synthetic image generation. ArXiv, arXiv:2304.09225v1. https://www.ncbi.nlm.nih.gov/pmc/articles/PMC10153296/

Hiersche, K., Schettini, E., Li, J., & Saygin, Z. (2024). Functional dissociation of the language network and other cognition in early childhood. Human Brain Mapping, 45(9), e26757.

Humphreys, G. F., Hoffman, P., Visser, M., Binney, R. J., & Lambon Ralph, M. A. (2015). Establishing task-and modality-dependent dissociations between the semantic and default mode networks. Proceedings of the National Academy of Sciences, 112(25), 7857–7862.

Isik, L., Koldewyn, K., Beeler, D., & Kanwisher, N. (2017). Perceiving social interactions in the posterior superior temporal sulcus. Proceedings of the National Academy of Sciences, 114(43), E9145–E9152. 10.1073/pnas.1714471114

Ivanova, A. A., Mineroff, Z., Zimmerer, V., Kanwisher, N., Varley, R., & Fedorenko, E. (2021). The Language Network Is Recruited but Not Required for Nonverbal Event Semantics. Neurobiology of Language, 2(2), 176–201. 10.1162/nol_a_00030

Ivanova, A. A., Srikant, S., Sueoka, Y., Kean, H. H., Dhamala, R., O’Reilly, U.-M., Bers, M. U., & Fedorenko, E. (2020). Comprehension of computer code relies primarily on domain-general executive brain regions. eLife, 9, e58906. 10.7554/eLife.58906

Janse, E., & Adank, P. (2012). Predicting foreign-accent adaptation in older adults. Quarterly Journal of Experimental Psychology (2006), 65(8), 1563–1585. 10.1080/17470218.2012.658822

Jouravlev, O., Schwartz, R., Ayyash, D., Mineroff, Z., Gibson, E., & Fedorenko, E. (2019). Tracking Colisteners’ Knowledge States During Language Comprehension. Psychological Science, 30(1), 3–19. 10.1177/0956797618807674

Kanwisher, N., McDermott, J., & Chun, M. M. (1997). The Fusiform Face Area: A Module in Human Extrastriate Cortex Specialized for Face Perception. Journal of Neuroscience, 17(11), 4302–4311. 10.1523/JNEUROSCI.17-11-04302.1997

Kauf, C., Kim, H. S., Lee, E. J., Jhingan, N., She, J. S., Taliaferro, M., Gibson, E., & Fedorenko, E. (2024). Linguistic inputs must be syntactically parsable to fully engage the language network (p. 2024.06.21.599332). bioRxiv. 10.1101/2024.06.21.599332

Kong, X.-Z., Tzourio-Mazoyer, N., Joliot, M., Fedorenko, E., Liu, J., Fisher, S. E., & Francks, C. (2020). Gene Expression Correlates of the Cortical Network Underlying Sentence Processing. Neurobiology of Language, 1(1), 77–103. 10.1162/nol_a_00004

Kuperberg, G. R., Sitnikova, T., Caplan, D., & Holcomb, P. J. (2003). Electrophysiological distinctions in processing conceptual relationships within simple sentences. Cognitive Brain Research, 17(1), 117–129. 10.1016/S0926-6410(03)00086-7

Kuznetsova, A., Brockhoff, P. B., & Christensen, R. H. B. (2017). lmerTest Package: Tests in Linear Mixed Effects Models. Journal of Statistical Software, 82, 1–26. 10.18637/jss.v082.i13

Landau, S. M., Schumacher, E. H., Garavan, H., Druzgal, T. J., & D’Esposito, M. (2004). A functional MRI study of the influence of practice on component processes of working memory. Neuroimage, 22(1), 211–221.

Lee, J. J., Scott, T. L., & Perrachione, T. K. (2024). Efficient functional localization of language regions in the brain. NeuroImage, 285, 120489. 10.1016/j.neuroimage.2023.120489

Leech, R., Kamourieh, S., Beckmann, C. F., & Sharp, D. J. (2011). Fractionating the default mode network: distinct contributions of the ventral and dorsal posterior cingulate cortex to cognitive control. Journal of Neuroscience, 31(9), 3217–3224.

Li, J., Hiersche, K. J., & Saygin, Z. M. (2024). Demystifying visual word form area visual and nonvisual response properties with precision fMRI. iScience, 27(12).

Lipkin, B., Tuckute, G., Affourtit, J., Small, H., Mineroff, Z., Kean, H., Jouravlev, O., Rakocevic, L., Pritchett, B., Siegelman, M., Hoeflin, C., Pongos, A., Blank, I. A., Struhl, M. K., Ivanova, A., Shannon, S., Sathe, A., Hoffmann, M., Nieto-Castañón, A., & Fedorenko, E. (2022). Probabilistic atlas for the language network based on precision fMRI data from >800 individuals. Scientific Data, 9(1), Article 1. 10.1038/s41597-022-01645-3

Liu, Y., Luo, C., Zheng, J., Liang, J., & Ding, N. (2022). Working memory asymmetrically modulates auditory and linguistic processing of speech. NeuroImage, 264, 119698. 10.1016/j.neuroimage.2022.119698

Liu, Y.-F., Kim, J., Wilson, C., & Bedny, M. (2020). Computer code comprehension shares neural resources with formal logical inference in the fronto-parietal network. eLife, 9, e59340. 10.7554/eLife.59340

Luria, A. R. (1970). The functional organization of the brain. Scientific American, 222(3), 66–72 passim. 10.1038/scientificamerican0370-66

MacGregor, L. J., Gilbert, R. A., Balewski, Z., Mitchell, D. J., Erzinçlioğlu, S. W., Rodd, J. M., Duncan, J., Fedorenko, E., & Davis, M. H. (2022). Causal Contributions of the Domain-General (Multiple Demand) and the Language-Selective Brain Networks to Perceptual and Semantic Challenges in Speech Comprehension. Neurobiology of Language, 3(4), 665–698. 10.1162/nol_a_00081

Mahowald, K., & Fedorenko, E. (2016). Reliable individual-level neural markers of high-level language processing: A necessary precursor for relating neural variability to behavioral and genetic variability. NeuroImage, 139, 74–93. 10.1016/j.neuroimage.2016.05.073

Malik-Moraleda, S., Ayyash, D., Gallée, J., Affourtit, J., Hoffmann, M., Mineroff, Z., Jouravlev, O., & Fedorenko, E. (2022). An investigation across 45 languages and 12 language families reveals a universal language network. Nature Neuroscience, 25(8), 1014–1019. 10.1038/s41593-022-01114-5

Malik-Moraleda, S., Jouravlev, O., Taliaferro, M., Mineroff, Z., Cucu, T., Mahowald, K., Blank, I. A., & Fedorenko, E. (2024). Functional characterization of the language network of polyglots and hyperpolyglots with precision fMRI. Cerebral Cortex, 34(3), bhae049. 10.1093/cercor/bhae049

Marvi, A. I., Hutchinson, S., Fedorenko, E., Saxe, R. R., Kamps, F. S., Regev, T. I., Chen, E.M., & Kanwisher, N. G. (2025). An efficient multifunction fMRI localizer for high-level visual, auditory, and cognitive regions in humans. Imaging Neuroscience, 3, IMAG-a.

Mattys, S. L., & Wiget, L. (2011). Effects of cognitive load on speech recognition. Journal of Memory and Language, 65(2), 145–160. 10.1016/j.jml.2011.04.004

Mineroff, Z., Blank, I. A., Mahowald, K., & Fedorenko, E. (2018). A robust dissociation among the language, multiple demand, and default mode networks: Evidence from inter-region correlations in effect size. Neuropsychologia, 119, 501–511. 10.1016/j.neuropsychologia.2018.09.011

Mollica, F., & Piantadosi, S. (2017). An incremental information-theoretic buffer supports sentence processing. Cognitive Science. https://www.semanticscholar.org/paper/An-incremental-information-theoretic-buffer-Mollica-Piantadosi/7def8be7453fad40fc41ce9d5f94125c7c5e0997

Mollica, F., Siegelman, M., Diachek, E., Piantadosi, S. T., Mineroff, Z., Futrell, R., Kean, H., Qian, P., & Fedorenko, E. (2020). Composition is the Core Driver of the Language-selective Network. Neurobiology of Language, 1(1), 104–134. 10.1162/nol_a_00005

Monti, M. M., Parsons, L. M., & Osherson, D. N. (2009). The boundaries of language and thought in deductive inference. Proceedings of the National Academy of Sciences of the United States of America, 106(30), 12554–12559. 10.1073/pnas.0902422106

Mur, M., Mitchell, D. J., Brüggemann, S., & Duncan, J. (2025). Stimulus effects dwarf task effects in human visual cortex. bioRxiv, doi: 10.1101/2025.06.18.660183

Naselaris, T., Allen, E., & Kay, K. (2021). Extensive sampling for complete models of individual brains. Current Opinion in Behavioral Sciences, 40, 45–51. 10.1016/j.cobeha.2020.12.008

Nieto-Castanon, A. (2020). Handbook of functional connectivity Magnetic Resonance Imaging methods in CONN. Hilbert Press.

Nieto-Castañón, A., & Fedorenko, E. (2012). Subject-specific functional localizers increase sensitivity and functional resolution of multi-subject analyses. NeuroImage, 63(3), 1646–1669. 10.1016/j.neuroimage.2012.06.065

Nieuwland, M. S., Martin, A. E., & Carreiras, M. (2012). Brain regions that process case: Evidence from basque. Human Brain Mapping, 33(11), 2509–2520. 10.1002/hbm.21377

Oldfield, R. C. (1971). The assessment and analysis of handedness: The Edinburgh inventory. Neuropsychologia, 9(1), 97–113. 10.1016/0028-3932(71)90067-4

Olson, H. A., Chen, E. M., Lydic, K. O., & Saxe, R. R. (2023). Left-Hemisphere Cortical Language Regions Respond Equally to Observed Dialogue and Monologue. Neurobiology of Language, 4(4), 575–610. 10.1162/nol_a_00123

Overath, T., McDermott, J. H., Zarate, J. M., & Poeppel, D. (2015). The cortical analysis of speech-specific temporal structure revealed by responses to sound quilts. Nature Neuroscience, 18(6), Article 6. 10.1038/nn.4021

Ozernov-Palchik, O., O’Brien, A. M., Lee, E. J., Richardson, H., Romeo, R., Lipkin, B., Small, H., Capella, J., Nieto-Castañón, A., Saxe, R., Gabrieli, J. D. E., & Fedorenko, E. (2024). Precision fMRI reveals that the language network exhibits adult-like left-hemispheric lateralization by 4 years of age (p. 2024.05.15.594172). bioRxiv. 10.1101/2024.05.15.594172

Potter, M. C. (2012). Recognition and Memory for Briefly Presented Scenes. Frontiers in Psychology, 3, 32. 10.3389/fpsyg.2012.00032

Potter, M. C., Kroll, J. F., & Harris, C. (1980). Comprehension and memory in rapid sequential reading. In Attention and Performance VIII (pp. 395–418). Hillsdale, NJ: Erlbaum.

Potter, M. C., Kroll, J. F., Yachzel, B., Carpenter, E., & Sherman, J. (1986). Pictures in sentences: Understanding without words. Journal of Experimental Psychology. General, 115(3), 281–294. 10.1037//0096-3445.115.3.281

Price, C. J. (2010). The anatomy of language: A review of 100 fMRI studies published in 2009. Annals of the New York Academy of Sciences, 1191, 62–88. 10.1111/j.1749-6632.2010.05444.x

Quillen, I. A., Yen, M., & Wilson, S. M. (2021). Distinct Neural Correlates of Linguistic and Non-Linguistic Demand. Neurobiology of Language, 2(2), 202–225. 10.1162/nol_a_00031

Ratan Murty, N. A., Bashivan, P., Abate, A., DiCarlo, J. J., & Kanwisher, N. (2021). Computational models of category-selective brain regions enable high-throughput tests of selectivity. Nature Communications, 12(1), Article 1. 10.1038/s41467-021-25409-6

Regev, T. I., Kim, H. S., Chen, X., Affourtit, J., Schipper, A. E., Bergen, L., Mahowald, K., & Fedorenko, E. (2024). High-level language brain regions process sublexical regularities. Cerebral Cortex, 34(3), bhae077. 10.1093/cercor/bhae077

Richardson, H., Koster-Hale, J., Caselli, N., Magid, R., Benedict, R., Olson, H., Pyers, J., & Saxe, R. (2020). Reduced neural selectivity for mental states in deaf children with delayed exposure to sign language. Nature Communications, 11(1), 3246. 10.1038/s41467-020-17004-y

Russ, B. E., Petkov, C. I., Kwok, S. C., Zhu, Q., Belin, P., Vanduffel, W., & Hamed, S. B. (2021). Common functional localizers to enhance NHP & cross-species neuroscience imaging research. NeuroImage, 237, 118203. 10.1016/j.neuroimage.2021.118203

Saxe, R., Brett, M., & Kanwisher, N. (2006). Divide and conquer: A defense of functional localizers. NeuroImage, 30(4), 1088–1096; discussion 1097-1099. 10.1016/j.neuroimage.2005.12.062

Saxe, R., & Kanwisher, N. (2003). People thinking about thinking people. The role of the temporo-parietal junction in “theory of mind.” NeuroImage, 19(4), 1835–1842. 10.1016/s1053-8119(03)00230-1

Schrimpf, M., Blank, I. A., Tuckute, G., Kauf, C., Hosseini, E. A., Kanwisher, N., Tenenbaum, J. B., & Fedorenko, E. (2021). The neural architecture of language: Integrative modeling converges on predictive processing. Proceedings of the National Academy of Sciences of the United States of America, 118(45), 1–12.

Scott, T. L., Gallée, J., & Fedorenko, E. (2017). A new fun and robust version of an fMRI localizer for the frontotemporal language system. Cognitive Neuroscience, 8(3), 167–176.

Shain, C., Blank, I. A., Fedorenko, E., Gibson, E., & Schuler, W. (2022). Robust Effects of Working Memory Demand during Naturalistic Language Comprehension in Language-Selective Cortex. The Journal of Neuroscience, 42(39), 7412–7430. 10.1523/JNEUROSCI.1894-21.2022

Shain, C., Blank, I. A., van Schijndel, M., Schuler, W., & Fedorenko, E. (2020). fMRI reveals language-specific predictive coding during naturalistic sentence comprehension. Neuropsychologia, 138, 107307. 10.1016/j.neuropsychologia.2019.107307

Shain, C., & Fedorenko, E. (2025). A language network in the individualized functional connectomes of over 1,000 human brains doing arbitrary tasks. bioRxiv, doi: 10.1101/2025.03.29.646067

Shain, C., Kean, H., Casto, C., Lipkin, B., Affourtit, J., Siegelman, M., Mollica, F., & Fedorenko, E. (2024). Distributed Sensitivity to Syntax and Semantics throughout the Language Network. Journal of Cognitive Neuroscience, 1–43. 10.1162/jocn_a_02164

Shain, C., Paunov, A., Chen, X., Lipkin, B., & Fedorenko, E. (2023). No evidence of theory of mind reasoning in the human language network. Cerebral Cortex, 33(10), 6299–6319. 10.1093/cercor/bhac505

Shashidhara, S., Mitchell, D. J., Erez, Y., & Duncan, J. (2019). Progressive Recruitment of the Frontoparietal Multiple-demand System with Increased Task Complexity, Time Pressure, and Reward. Journal of Cognitive Neuroscience, 31(11), 1617–1630. 10.1162/jocn_a_01440

Shashidhara, S., Spronkers, F. S., & Erez, Y. (2020). Individual-subject functional localization increases univariate activation but not multivariate pattern discriminability in the ‘multiple-demand’ frontoparietal network. Journal of Cognitive Neuroscience, 32(7), 1348–1368. 10.1162/jocn_a_01554

Somers, D. C., Michalka, S. W., Tobyne, S. M., & Noyce, A. L. (2021). Individual subject approaches to mapping sensory-biased and multiple-demand regions in human frontal cortex. Current Opinion in Behavioral Sciences, 40, 169–177. 10.1016/j.cobeha.2021.05.002

Tahmasebi, A. M., Artiges, E., Banaschewski, T., Barker, G. J., Bruehl, R., Büchel, C., Conrod, P. J., Flor, H., Garavan, H., Gallinat, J., Heinz, A., Ittermann, B., Loth, E., Mareckova, K., Martinot, J.-L., Poline, J.-B., Rietschel, M., Smolka, M. N., Ströhle, A., … Consortium, T. I. (2012). Creating probabilistic maps of the face network in the adolescent brain: A multicentre functional MRI study. Human Brain Mapping, 33(4), 938–957. 10.1002/hbm.21261

Terhune-Cotter, B., McCullough, S., & Emmorey, K. (2023). Functional Localizers for American Sign Language Comprehension. Society for the Neurobiology of Language. https://www.neurolang.org/2023/presentation/

Tomaiuolo, F., MacDonald, J. D., Caramanos, Z., Posner, G., Chiavaras, M., Evans, A. C., & Petrides, M. (1999). Morphology, morphometry and probability mapping of the pars opercularis of the inferior frontal gyrus: An in vivo MRI analysis. The European Journal of Neuroscience, 11(9), 3033–3046. 10.1046/j.1460-9568.1999.00718.x

Tuckute, G., Kanwisher, N., & Fedorenko, E. (2024a). Language in brains, minds, and machines. Annual Review of Neuroscience, 47(2024), 277–301.

Tuckute, G., Sathe, A., Srikant, S., Taliaferro, M., Wang, M., Schrimpf, M., Kay, K., & Fedorenko, E. (2024b). Driving and suppressing the human language network using large language models. Nature Human Behaviour, 8, 544–561. 10.1038/s41562-023-01783-7

Tuckute, G., Lee, E. J., Ou, Y., Fedorenko, E., & Kay, K. (2025). A two-dimensional space of linguistic representations shared across individuals (p. 2025.05.21.655330). bioRxiv, doi: 10.1101/2025.05.21.655330

Vagharchakian, L., Dehaene-Lambertz, G., Pallier, C., & Dehaene, S. (2012). A Temporal Bottleneck in the Language Comprehension Network. Journal of Neuroscience, 32(26), 9089–9102. 10.1523/JNEUROSCI.5685-11.2012

Vázquez-Rodríguez, B., Suárez, L. E., Markello, R. D., Shafiei, G., Paquola, C., Hagmann, P., van den Heuvel, M. P., Bernhardt, B. C., Spreng, R. N., & Misic, B. (2019). Gradients of structure–function tethering across neocortex. Proceedings of the National Academy of Sciences, 116(42), 21219–21227. 10.1073/pnas.1903403116

Wang, B., LeBel, A., & D’Mello, A. M. (2025). Ignoring the cerebellum is hindering progress in neuroscience. Trends in Cognitive Sciences, 29(4), 318–330

Wehbe, L., Blank, I. A., Shain, C., Futrell, R., Levy, R., von der Malsburg, T., Smith, N., Gibson, E., & Fedorenko, E. (2021). Incremental Language Comprehension Difficulty Predicts Activity in the Language Network but Not the Multiple Demand Network. Cerebral Cortex (New York, N.Y.: 1991), 31(9), 4006–4023. 10.1093/cercor/bhab065

Wilson, S. M., Entrup, J. L., Schneck, S. M., Onuscheck, C. F., Levy, D. F., Rahman, M., Willey, E., Casilio, M., Yen, M., Brito, A. C., Kam, W., Davis, L. T., de Riesthal, M., & Kirshner, H. S. (2023). Recovery from aphasia in the first year after stroke. Brain, 146(3), 1021–1039. 10.1093/brain/awac129

Wolna, A., Szewczyk, J., Diaz, M., Domagalik, A., Szwed, M., & Wodniecka, Z. (2024). Domain-general and language-specific contributions to speech production in a second language: An fMRI study using functional localizers. Scientific Reports, 14(1), 57. 10.1038/s41598-023-49375-9

Wolna, A., Wright, A., Casto, C., Lipkin, B., & Fedorenko, E. (2025). The extended language network: Language selective brain areas whose contributions to language remain to be discovered. bioRxiv, doi: 10.1101/2025.04.02.646835

Xiao, W., & Kreiman, G. (2020). XDream: Finding preferred stimuli for visual neurons using generative networks and gradient-free optimization. PLOS Computational Biology, 16(6), e1007973. 10.1371/journal.pcbi.1007973

Yamins, D. L. K., Hong, H., Cadieu, C. F., Solomon, E. A., Seibert, D., & DiCarlo, J. J. (2014). Performance-optimized hierarchical models predict neural responses in higher visual cortex. Proceedings of the National Academy of Sciences, 111(23), 8619–8624. 10.1073/pnas.1403112111

